# Protein kinase C is a key target for attenuation of Leigh syndrome by rapamycin

**DOI:** 10.1101/562207

**Authors:** Miguel Martin-Perez, Takashi K. Ito, Anthony S. Grillo, Anthony S. Valente, Jeehae Han, Samuel Entwisle, Heather Z. Huang, Dayae Kim, Masanao Yajima, Matt Kaeberlein, Judit Villén

## Abstract

Leigh syndrome is a fatal neurometabolic disorder caused by defects in mitochondrial function. mTOR inhibition with rapamycin attenuates disease progression in a mouse model of Leigh syndrome (Ndufs4 KO mouse); however, the mechanism of rescue is unknown. Here we assessed the impact of rapamycin on the brain proteome and phosphoproteome of Ndufs4 KO mice. We report that rapamycin remodels the brain proteome to alter mitochondrial structure, inhibits signaling through both mTOR complexes, and inhibits multiple protein kinase C (PKC) isoforms. Administration of PKC inhibitors was sufficient to increase survival, delay neurological deficits, and prevent hair loss in Ndufs4 KO mice. Thus, PKC may be a viable therapeutic target for treating severe mitochondrial disease.

**One Sentence Summary:** Proteomic and phosphoproteomic analysis of mouse brain identifies PKC as a key target to treat mitochondrial disease

## Main Text

Mitochondria are essential organelles, generating 80-90% of cellular ATP via respiratory metabolism (*1*). Genetic impairment of the mitochondrial respiratory chain can lead to several mitochondrial diseases with diverse clinical etiology, but often associated with encephalopathy, neuroinflammation, and neurodegeneration. Leigh syndrome is regarded as the most common childhood mitochondrial disease and is characterized by bilateral lesions in the brainstem and basal ganglia. Mutations in both nuclear and mitochondrially encoded components of the electron transport chain, such as Ndufs4 of Complex I (C-I), have been identified as causal for Leigh Syndrome (*2*). Knockout of the Ndufs4 gene in mice (Ndufs4 KO) leads to rapid onset of a mitochondrial disease that shares many clinical features of Leigh Syndrome including neurodegeneration, neuroinflammation, progressive lesions in the brain, and substantially shortened lifespan (*3*).

The mechanistic target of rapamycin (mTOR) is a serine/threonine kinase that controls cell growth and metabolism and is hyperactivated in several animal models of mitochondrial disease (*4*-*6*). mTOR functions in two distinct complexes: mTOR complex 1 (mTORC1) and mTOR complex 2 (mTORC2). We previously reported that the mTORC1 inhibitor rapamycin alleviates neuropathic symptoms in Ndufs4 KO mice and nearly triples their life expectancy (*4*). Rapamycin treatment also induces a profound metabolic shift in this context, reducing accumulation of glycolytic intermediates and lactic acid, and suppressing the extremely low body fat of Ndufs4 KO mice (*4*).

To further explore the mechanisms of rapamycin-mediated rescue in Ndufs4 KO mice, we used mass spectrometry-based proteomics to elucidate changes in the brain proteome and phosphoproteome. We analyzed brain tissue of 30-day old wild-type and Ndufs4 KO mice treated with a daily intraperitoneal injection of vehicle or 8 mg/kg/day rapamycin for 20 days (Fig.1A). Postnatal day 30 is approximately one week before the appearance of neurodegeneration and behavioral abnormalities in Ndufs4 KO mice (*3, 4*). As expected (*4*), rapamycin-treated Ndufs4 KO mice had lower body weight and brain weight relative to vehicle-treated mice (Fig. 1B).

**Figure 1.**
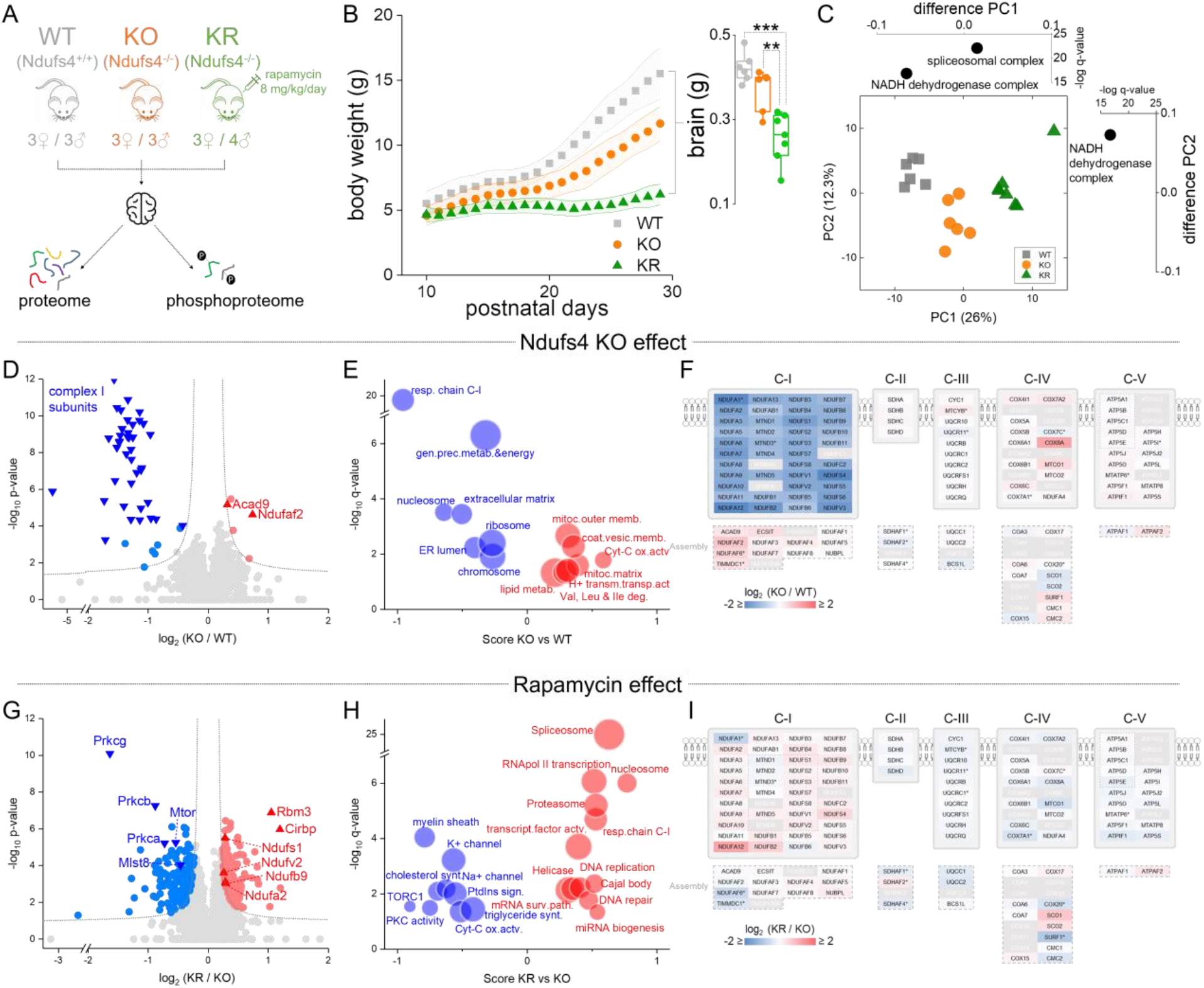
Rapamycin remodels the brain proteome in Ndufs4 deficient mice. (**A**) Experimental design. (**B**) Mice body weight from the three experimental groups (average ± s.d.). Inner graphs show total brain weight at the end of the experimental trial (30 days) (t-test: ** p < 0.01; *** p < 0.001). (**C**) PCA analysis of log_2_ transformed normalized protein abundance data. Side graphs indicate significantly enriched GO slim terms in the loadings for each component. (**D,E**) Volcano plots comparing protein abundance in Ndufs4 KO (KO) *vs.* wild type (WT) groups at the level of individual proteins (D) or representative biological terms (E). (**F**) Mapping of protein abundance differences in individual subunits of the respiratory chain between the Ndufs4 KO and WT groups. Assembly subunits of each mitochondrial respiratory complex are displayed at the bottom of each plot. Normalized abundance data was used, except for subunits with asterisk where intensity raw values were used. In grey are subunits with no abundance information. (**G**, **H**) Volcano plots comparing protein abundance in rapamycin-treated Ndufs4 KO (KR) *vs.* KO groups at the level of individual proteins (G) or representative biological terms (H). In (D) and (G), dotted lines indicate cut-off for significant changes: t-test FDR q-value < 0.05 and artificial within groups variance S0 = 0.1). (E) and (H) show significantly enriched annotation terms from 1D enrichment analysis of log_2_ differences between groups; dot size is proportional to the number of proteins within that term. (**I**) Same as (F) but between Ndufs4 KO with (KR) and without (KO) rapamycin groups.

We quantified 6,231 proteins (Data S1) with high reproducibility (Fig. S1A), of which 16% (1,004 proteins) changed significantly in abundance among groups (ANOVA FDR q-value < 0.05). The mitochondrial proteome showed substantial changes in response to loss of Ndufs4, driven primarily by altered abundance of C-I proteins (Fig. S1A). In agreement, principal component analysis (PCA) revealed changes to C-I proteins as the major discriminant among groups (Fig. 1C). Most C-I proteins decreased substantially in Ndufs4 KO mouse brains (Fig. 1D, 1E, 1F, Fig. S2), except for two C-I assembly proteins (Acad9 and Ndufaf2) (Fig. 1D, 1F). Rapamycin treatment caused a modest increase of several C-I subunits (Fig. 1G, 1H, 1I, Fig. S2), but is not sufficient to restore C-I function (*4*). In addition to the expected decrease in C-I abundance in Ndufs4 KO brain, we observed changes in other respiratory chain protein complexes (Fig. 1F, 1I, Fig. S2). In particular, Ndufs4 KO mice showed an increase in the cytochrome c oxidase complex (C-IV or COX), which was reverted by rapamycin treatment.

Rapamycin treatment of Ndufs4 KO mice caused several additional changes in the brain proteome. Most notably, rapamycin significantly decreased mTOR and mLST8 (Fig. 1G), which are both core components of mTORC1 and mTORC2. Although rapamycin is a specific inhibitor of mTORC1 (*7, 8*), chronic rapamycin treatment has been reported to inhibit mTORC2 in mice (*9, 10*), and our data indicates that this may be mediated by decreased abundance of mTOR and mLST8. In agreement with mTORC2 inhibition, we observed a reduction of phosphorylation at S473 of Akt (Fig. S3) and of protein kinase C (PKC), known mTORC2 targets (*11, 12*). All conventional PKC isoforms (PKC-α, PKC-β, and PKC-γ) were significantly decreased (Fig. 1G and Fig. S3B). Rapamycin treatment also significantly increased the abundance of the cold-inducible RNA-binding proteins Cirbp and Rbm3 (Fig. 1G), which have been reported to have neuroprotective effects (*13, 14*). Rapamycin also increased histone abundance and up-regulated processes associated with mTOR inhibition like mRNA splicing and proteasomal degradation, while down-regulating others such as myelination (Fig. 1H).

Hierarchical clustering of protein abundances grouped proteins into 4 clusters according to changes among the different groups (Fig. 2A). Clusters 1 and 4 contain protein changes associated with rapamycin treatment, while cluster 3 includes protein changes associated with genotype. Cluster 2 was composed of proteins that increased in Ndufs4 KO mice and restored to normal levels with rapamycin, and was significantly enriched in mitochondrial proteins. Among these, we found Hk1, the brain-specific hexokinase responsible for glucose utilization, which mostly localizes to the outer mitochondrial membrane (OMM) to promote the energetically favorable coupling of glycolysis to oxidative phosphorylation (*15*) (Fig. 2B). This agrees with our prior observations indicating that rapamycin shifts metabolism away from glycolysis in Ndufs4 KO mice (*4*). We found other OMM proteins significantly enriched in cluster 2, especially proteins involved in mitochondrial fission (Fig. 2C, 2D). Among proteins which decreased in KO mice and returned to normal levels with rapamycin we found Lppr1 (Fig. 2B), a neuronal growth promoter (*16*). We also found that increased levels of Mpst in KO mice, a neuroprotectant mitochondrial enzyme involved in hydrogen sulfide production in the brain, were further enhanced by rapamycin (Fig. 2B).

**Figure 2.**
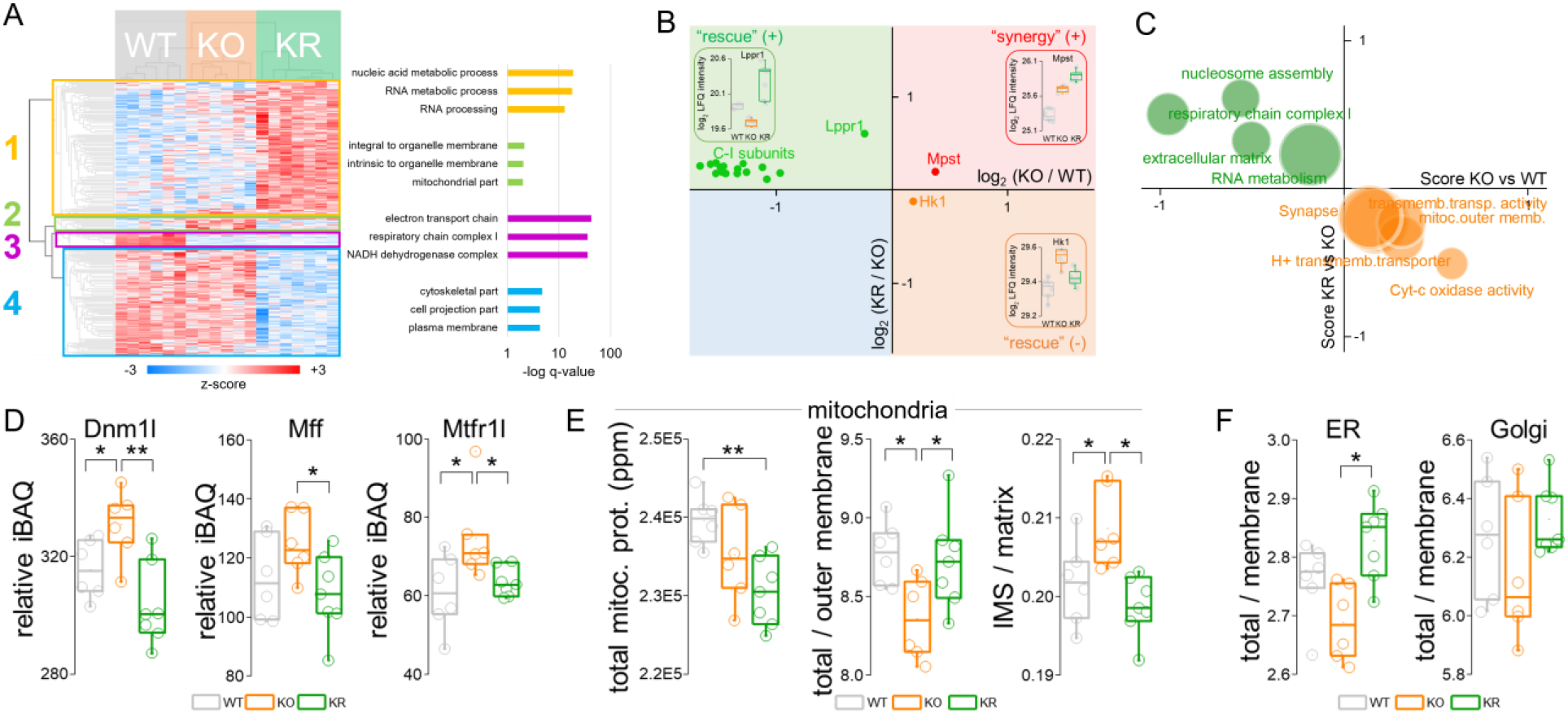
Rapamycin restores normal mitochondrial structure and functionality in Ndufs4 KO mice. (**A**) Heat map and hierarchical clustering of proteins with significant changes in abundance (ANOVA FDR q-value < 0.05). Top 3 significantly enriched GO terms from main clusters are shown on the right panel. (**B**, **C**) Scatter plot of individual protein abundances (B) and GO terms (C) showing significant changes (FDR q-value < 0.05) between KO *vs.* WT and KR *vs.* KO groups, in the same direction (i.e. rapamycin enhances Ndufs4 KO changes) or in opposite directions (i.e. rapamycin rescues Ndufs4 KO changes). (**D**) Changes in abundance of proteins involved in mitochondrial fission (iBAQ = intensity based absolute quantification). (**E**) Global changes in mitochondrial proteins. We calculate the ratio of total mitochondria:outer mitochondrial membrane protein as a proxy for the volume to surface area to monitor changes in organelle size (or shape), while the ratio mitochondrial intermembrane space (IMS):matrix protein tracks changes to cristae volume. (**F**) Changes in the ratio of total:organelle membrane protein for endoplasmic reticulum (ER) and Golgi apparatus, as a proxy for organelle size. (t-test: * p < 0.05; ** p < 0.01).

To gain information about potential changes in mitochondrial morphology (e.g. number, size, and shape), we calculated abundance ratios between different submitochondrial localizations. As a proxy for the volume to surface area to monitor changes in organelle size (or shape), we calculated the ratio of total mitochondrial protein abundances to OMM protein abundances (see methods, Fig. S4). Based on this analysis, brain mitochondria appeared to shrink in Ndufs4 KO mice (Fig. 2E), which agrees with the fragmentation of the mitochondrial network observed in cells derived from the same mouse model (*17*) and Leigh syndrome patients (*18, 19*). Furthermore, brains from Ndufs4 KO mice had lower matrix-to-intermembrane space protein abundance ratio than wild-type brains, perhaps indicative of reduced mitochondrial matrix volume (Fig. 2D). This coincides with the abnormally swollen cristae described in mitochondria from Ndufs4 KO mouse brain (*3*) and Leigh syndrome patients (*20*). Interestingly, similar calculation of the ratio between organelle membrane and total organelle protein abundances suggests that the endoplasmic reticulum (ER) also has reduced volume in knockout mouse brain (Fig. 2F), while other organelles (e.g. Golgi apparatus) remained unchanged (Fig. 2F). These apparent alterations in mitochondrial and ER structure were suppressed by rapamycin treatment.

Given that mTOR functions as a kinase, we also sought to assess changes in protein phosphorylation upon treatment with rapamycin. We quantified 15,971 phosphorylation sites (Data S2) in 3,091 proteins with high reproducibility (Fig. S5); 26% of these sites have not been previously reported. Globally, the phosphoproteome was altered to a lesser extent than the proteome (10% phosphosites with significant changes; ANOVA FDR q-value < 0.05), and most of the changes were associated with rapamycin treatment (Fig. 3A, 3C). Ndufs4 KO induced few significant changes in brain protein phosphorylation, consistent with the minimal changes in kinase abundance (Fig. 3B, 3D). Of all the phosphorylation sites detected by our analysis, only 2% belong to mitochondrial proteins, and a very few changed significantly after rapamycin treatment (Fig. S5). These involve proteins related to mitochondrial transport, metabolism and fission (Table S1). The low occurrence of phosphorylation events on mitochondrial proteins suggests that mitochondrial function is primarily regulated by changes in protein and substrate abundance or possibly by other post-translational modifications.

**Figure 3.**
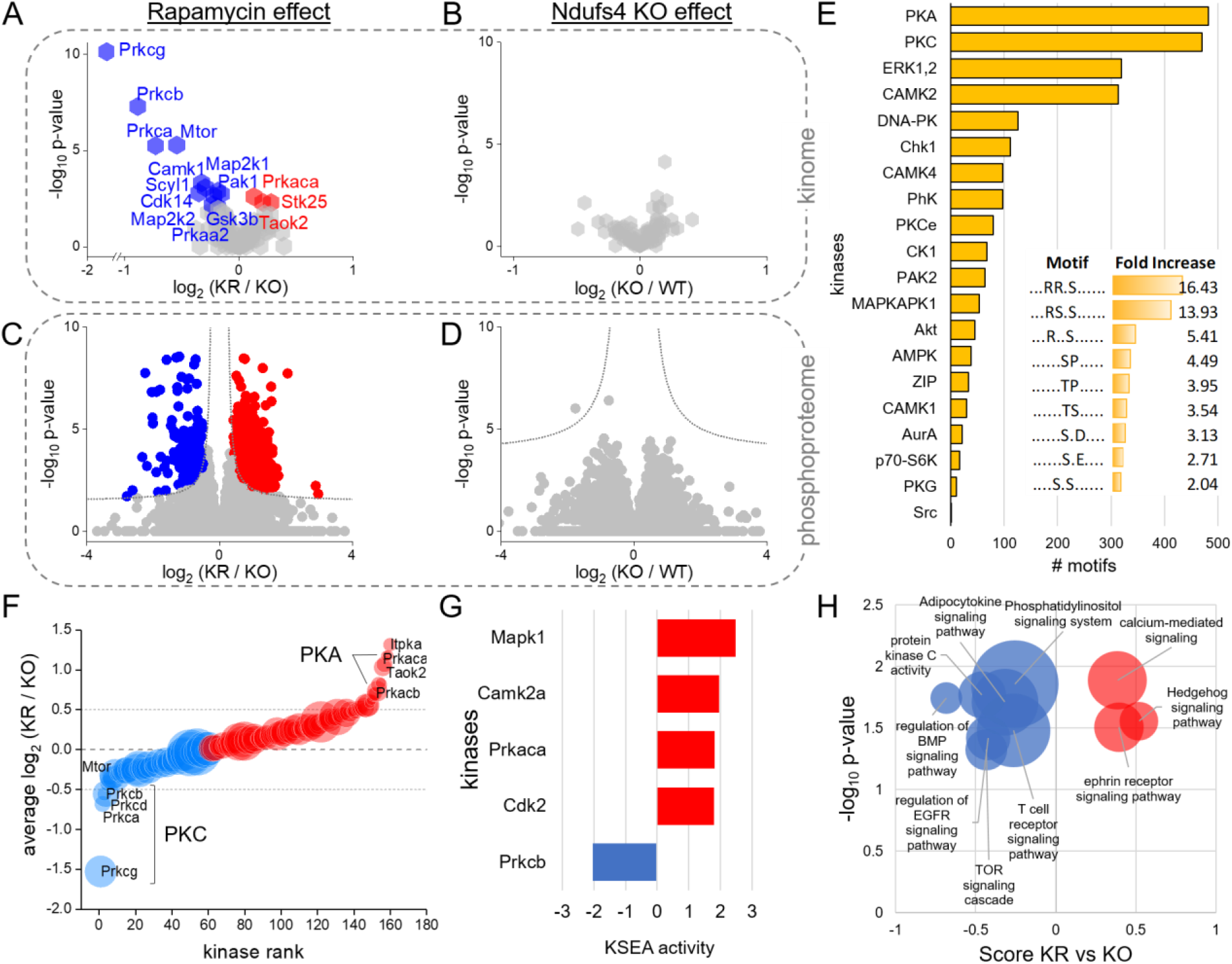
Rapamycin reduces PKC activity in Ndufs4 KO mouse brain. (**A**, **B**) Volcano plots comparing the abundance of protein kinases between KR and KO (rapamycin effect) or KO and WT (genotype effect) groups. Kinases with significant changes are colored and labeled (t-test FDR q-value < 0.05) (**C**, **D**) Volcano plots comparing phosphosite abundance differences in KR *vs.* KO (rapamycin effect) and KO *vs.* WT (genotype effect) groups. Dotted lines indicate cut-off for significant changes: t-test FDR q-value < 0.05 and artificial within groups variance S0 = 0.1). (**E**) Enriched kinase motifs in significantly changing phosphosites (Fisher exact test FDR q-value < 0.05). Inner graph shows overrepresented linear sequence motifs. (**F**) Average rapamycin-induced changes in phosphorylation of kinases. Dot size is proportional to the number of phosphosites considered for each kinase. Only shown here kinases with n≥3 sites. (**G**) Kinase Substrate Enrichment Analysis (KSEA) showing kinases with significant changes in activity upon rapamycin treatment (p-value < 0.05). (**H**) Volcano plot showing signaling pathways from 1D enrichment analysis of phosphorylation changes between KR and KO. Relative enrichment at the protein level was used to prevent overrepresentation of multiple sites from the same protein. Dot size is proportional to the number of proteins for each annotated term.

We detected several changes in protein phosphorylation with rapamycin treatment in Ndufs4 KO mice, including several mTOR complex members and known mTOR substrates (Fig. S6). Interestingly, global changes in protein phosphorylation showed significant correlation with changes to protein abundance (Fig. S7), indicating that twenty days of rapamycin treatment largely rewires the signaling network in mouse brain by changing protein levels. Similar effects were observed in wild type and Ndufs4 KO mice (Supplementary text, Fig. S8). To determine which kinases target these phosphosites, we identified linear sequence motifs that were enriched in the set of rapamycin-regulated phosphosites (Fisher-test FDR q <0.05). We found an overrepresentation of basophilic motifs (e.g. RRxS*; RSxS*; S* = phosphoserine, x=any residue), which are typically targeted by kinases in the AGC (PKC and PKA) and CamK (CAMK2) families (Fig. 3E). Protein kinase C (PKC) and protein kinase A (PKA) themselves also exhibited significant changes upon rapamycin treatment, with PKC decreasing and PKA increasing in protein abundance (Fig. 3A) and phosphorylation levels (Fig. 3F). To assess changes in kinase activity, we applied kinase-substrate enrichment analysis (KSEA) (*21*), revealing that chronic treatment with rapamycin repressed PKC-β and induced PKA and CAMK2 activity (Fig. 3G). This result was further supported by direct inspection of phosphorylation changes of known substrates of these kinases (Fig. S9) and the kinase activation loop sites (Fig. S10). Changes in the activity of these kinases, also manifested in a tighter control of calcium homeostasis upon rapamycin treatment (Supplementary text, Fig. S11).

Among the multiple PKC isoforms showing reduced abundance upon rapamycin treatment, PKC-β appeared to show the most significant decrease in activity (Fig. 3G). PKC-β mediates activation of the NF-κB pathway via IKK-α and IκB phosphorylation and plays a key role in the activation of the immune system and inflammatory response (*22*). In addition to the inhibition of PKC-β with rapamycin treatment, we observed a global decrease in phosphorylation of proteins from pro-inflammatory signaling pathways (e.g. adipocytokine signaling system, T cell receptor signaling pathway, BMP signaling pathway) (Fig. 3H) and a decrease in abundance (Fig. 3A) together with an increase of inhibitory phosphorylation (Fig. S10) of the inflammation regulator Gsk3b (*23*). Western blot analysis of brain lysates confirmed downregulation of PKC-β phosphorylation (Fig. S3) as well as phosphorylation of IKKα (Fig. S12). Consistent with the role of IKKα in activating NF-κB by phosphorylating the NF-κB suppressor IκB and promoting its degradation, we observed rapamycin treatment decreased IκB phosphorylation while increasing total IκB levels compared to vehicle-treated mice (Fig. S12).

Based on the proteomic and phosphoproteomic data, we hypothesized that chronic rapamycin treatment attenuates inflammation and neurodegeneration in Ndufs4 KO mice in part through decreasing the activity of PKC-β. To directly test this, we treated Ndufs4 KO mice with three different PKC inhibitors: the pan-PKC inhibitors GO6983 and GF109203X, and the PKC-β specific inhibitor ruboxistaurin. All three drugs were able to significantly delay the onset of neurological symptoms (i.e. clasping) and increase survival of Ndufs4 KO mice (Fig. 4A and 4B). In contrast to rapamycin, however, the PKC inhibitors had no significant effects on growth and weight gain (Fig. 4C). We also observed that all three PKC inhibitors largely suppress the hair loss phenotype displayed by Ndufs4 KO mice at weaning (Fig. 4D and Fig. S13), which has been attributed to an NF-κB mediated inflammatory response in skin (*24*).

**Figure 4.**
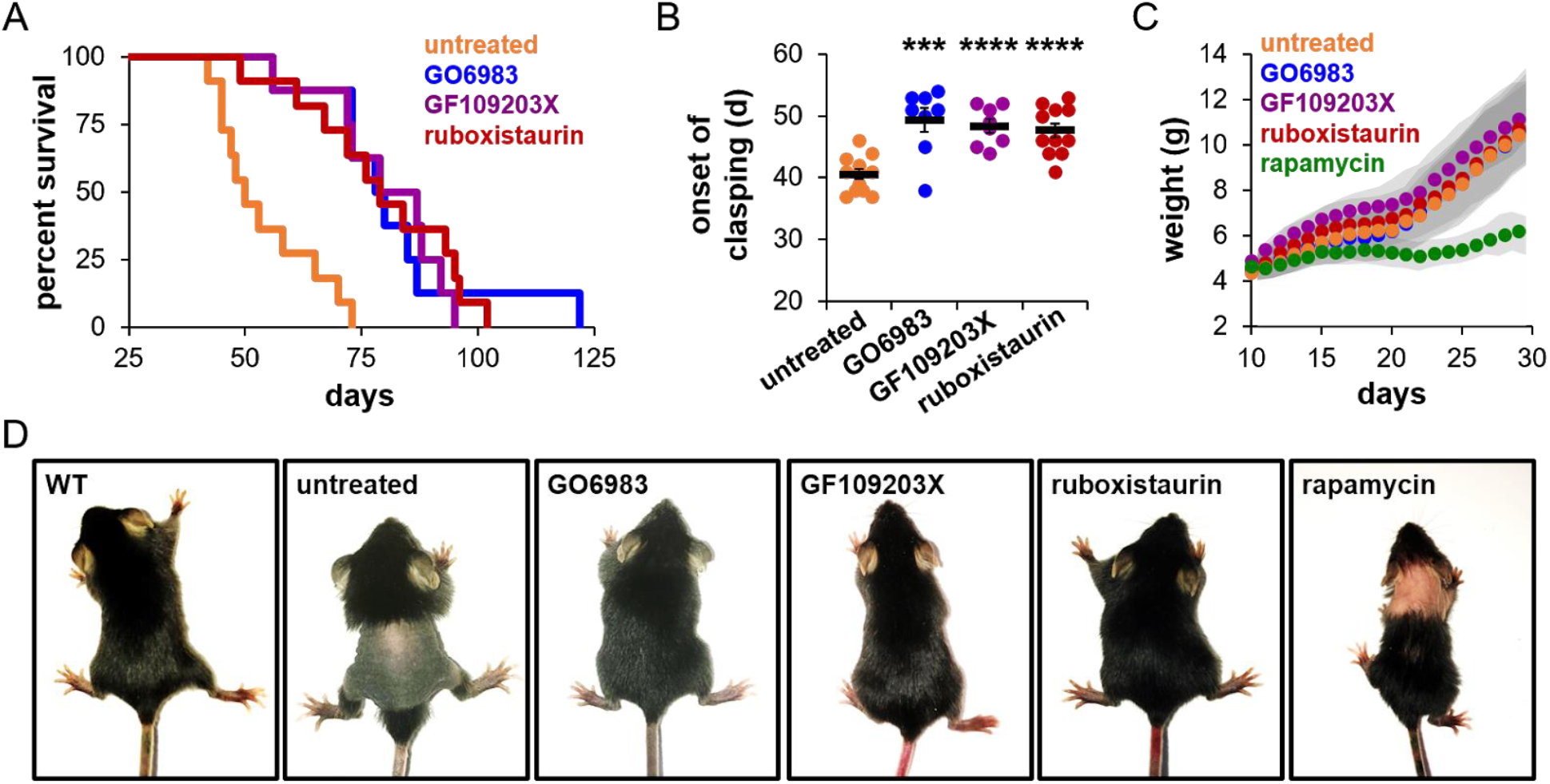
Inhibition of PKC signaling improves survival of mice with complex I deficiency. (**A**) Survival of Ndufs4 KO mice treated with pan-PKC inhibitors GO6983 (N = 8) and GF109203X (N = 8), the PKC-β inhibitor ruboxistaurin (N = 11), and their untreated control (N = 12). Untreated vs GO6983 p = 0.0003. Untreated vs GF109203X p = 0.0004, Untreated vs. ruboxistaurin p = 0.0003, log-rank test. (**B**) Day of starting clasping phenotype as a sign of neurodegeneration. *** p<0.001; **** p<0.0001. (**C**) Weights of Ndufs4 KO mice treated with vehicle, PKC inhibitors, or rapamycin. (**D**) PKC inhibitors largely prevent the alopecia phenotype of untreated Ndufs4 KO mice normally observed at weaning (representative images on postnatal day 21).

Taken together, these results suggest that inhibition of PKC-β contributes to suppression of mitochondrial disease and increased survival from chronic rapamycin treatment in Ndufs4 KO mice. These effects may be mediated through reduced abundance and activity of mTORC2, a known regulator of PKC. It has been suggested that inhibition of mTORC2 upon chronic rapamycin treatment leads to metabolic dysregulation and detrimental side effects in the context of normative aging in mice (*25, 26*); however, our data raise the possibility that inhibition of mTORC2 may also have beneficial anti-inflammatory and metabolic consequences, at least in the context of severe mitochondrial disease. Recently, it was reported that rapamycin and rapamycin-derivatives can improve outcomes in patients suffering from MELAS (Mitochondrial Encephalopathy, Lactic acidosis, and Stroke-like episodes) Syndrome (*27*), an adult-onset mitochondrial disease. The findings presented here suggest that PKC inhibitors may also have therapeutic value for treating mitochondrial disease and may be particularly useful in patients where side effects associated with mTOR inhibition represent a significant concern.

## Acknowledgments

We thank Victor V. Pineda, Nicolas J. LeTexier, James Phillips, Jo Tan, Yenna Lee, Tuyet Nguyen, Samy Khessib, Natalie Lim, Chenhao Lu, Surapat Mekvanich, and Azaad O. Zimmermann for assisting with animal experiments. **Funding:** This work was supported by NIH grants R01 NS098329 and R01 AG056359 (to M.K. and J.V.), and R35 GM119536 (to J.V.). T.K.I. was supported by a JSPS Postdoctoral Fellowship and a Uehara Memorial Foundation Postdoctoral Research Fellowship. A.S.G. was supported by a Ruth L. Kirschstein NRSA through the NIH NINDS (F32NS110109). **Author contributions:** M.M.-P. designed, conducted, analyzed, and interpreted all the proteomic experiments; developed methodology to assess cell morphological changes using proteomic data; prepared most of the figures; and wrote the initial draft of the paper. T.K.I. conceptualized the study; designed, conducted, and interpreted phenotypic and lifespan experiments with rapamycin and broad-spectrum PKC inhibitors; and revised and edited the manuscript. A.S.G. designed, conducted, and interpreted phenotypic and lifespan experiments with the PKC-beta inhibitor; conducted and interpreted all the Western blots, preparing associated figures; and revised and edited the manuscript. A.V. conducted KSEA analysis and revised and edited the manuscript. J.H. conceptualized the study and obtained the brain tissue samples for proteomic analysis. S.E. assisted with the proteomic analysis and revised and edited the manuscript. H.Z.H. and D.K. assisted with mouse experiments. M.Y. assisted with statistical analysis and revised and edited the manuscript. M.K. conceived and coordinated the project; supervised the mouse work; provided animal resources and funding for T.K.I. A.S.G., J.H., H.Z.H., and D.K.; and revised and edited the manuscript. J.V. coordinated the project; supervised the proteomics work; provided instrumentation resources and funding for M.M.P., A.V. and S.E.; and revised and edited the manuscript. **Competing interests:** The authors declare no competing interests. **Data and Materials availability:** All mass spectrometry raw files and searches have been deposited in the MassIVE repository with dataset identifier PXD012158. Protein and phosphorylation quantification results are provided as Supplementary Data.

## Supplementary Materials

### Materials and Methods

#### Animals and animal care

All studies with mice were performed in strict accordance with the Guide for the Care and Use of Laboratory Animals of the National Institutes of Health. The protocols used were approved by the University of Washington Institutional Animal Care and Use Committee. Mice were provided food, water, and/or gel *ad libitum* and were maintained on a 12:12-hr light/dark cycle.

*Ndufs4*^*+/-*^ mice were generously provided by the Palmiter Laboratory at the University of Washington and backcrossed with C57Bl/6 CR mice. Breeders of heterozygous *Ndufs4*^*+/-*^ mice were used and offspring were genotyped by PCR to identify *Ndufs4*^*+/+*^, *Ndufs4*^*+/-*^, and *Ndufs4*^*-/-*^ pups. Mice were weaned at 21-28 days of age. *Ndufs4*^*-/-*^ mice used in lifespan experiments were housed with one control littermate (*Ndufs4*^*+/+*^ or *Ndufs4*^*+/-*^) for warmth and companionship.

*Ndufs4*^*-/-*^ mice used in proteomic experiments were housed with one *Ndufs4*^*+/+*^ control littermate for warmth and companionship. Both male and female mice were used for all experiments, with no discernible differences observed in these experiments. Mice were weighed daily, and food and gel were provided on the bottom of cage. Mice used in lifespan experiments were checked daily for signs of clasping and were euthanized if they showed a 30% loss in maximum body weight or loss of righting reflex, were found prostrate, or were generally unresponsive.

#### Drug treatment

Rapamycin (LC Laboratories) and GO6983 (LC Laboratories) were dissolved in DMSO and diluted 100-fold in 5% PEG-400/5% Tween-80 (vehicle), sterile filtered, aliquoted into 1 mL portions, and stored at −80 °C until needed for use. GF109203X (Cayman Chemical) and ruboxistaurin hydrochloride (Synnovator) were directly dissolved in vehicle containing 1% DMSO. Starting at postnatal day 10 (P10), mice were treated daily between 2:00-6:00 p.m. (via intraperitoneal injection with 29 ½ gauge, 300 µL syringes) using 6.66 µL/g body weight of these solutions for final doses of 10 mg/kg/day (ruboxistaurin), 8 mg/kg/day (rapamycin and GF109203X), or 4 mg/kg/day (GO6983). Control mice were generally treated with vehicle using 6.66 µL/g body weight. For the lifespan experiments control mice were untreated, as it has been reported that untreated *Ndufs4*^*-/-*^ mice exhibit identical phenotypes as vehicle-treated *Ndufs4*^*-/-*^ mice (*4*), and we also show this here for lifespan, onset of clasping and weight gain (Fig. S14).

#### Sample preparation for proteomic analysis

WT and *Ndufs4*^*-/-*^ mice were treated daily with rapamycin or vehicle as described above from P10 until P29. At that age, mice were fasted overnight (for 12 h), treated with rapamycin, and re-fed in the morning for 4.5 hours. Unanesthetized mice were then euthanized by cervical dislocation. Brains were immediately isolated, flash frozen in liquid N_2_ and stored at −80°C. Frozen brains were first weighed and then grinded in liquid nitrogen. About 15 mg of ground tissue were resuspended in 600 μL of lysis buffer composed of 8M urea, 75mM NaCl, 50 mM Tris pH 8.2, and a mix of protease inhibitors (Roche Complete EDTA-free) and phosphatase inhibitors (50 mM beta-glycerophosphate, 10 mM sodium pyrophosphate, 1 mM sodium orthovanadate and 50 mM sodium fluoride). Samples were then subjected to 2 cycles of bead beating (1 min beating, 1.5 min rest) with 0.5mm diameter zirconia beads and sonicated for 5 min in ice. Samples were centrifuged at 4°C to remove debris and lysate protein concentration was measured by BCA assay (Pierce). Protein was reduced with 5 mM dithiothreitol (DTT) for 30 min at 55°C and alkylated with 15 mM iodoacetamide in the dark for 30 min at room temperature. The alkylation reaction was quenched by incubating with additional 5 mM DTT for 15 min at room temperature. Samples were diluted five-fold with 50 mM Tris pH 8.2. Proteolytic digestion was performed by adding trypsin at 1:200 enzyme to protein ratio and incubating at 37°C overnight. The digestion was quenched by addition of trifluoroacetic acid to pH 2. Samples were centrifuged to remove insoluble material and peptides were desalted over a 50 mg tC18 SepPak cartridge (Waters). Briefly, cartridges were conditioned with 1 mL of methanol, 3 mL of 100% acetonitrile, 1 mL of 70% acetonitrile, 0.25% acetic acid and 1 mL of 40% acetonitrile, 0.5% acetic acid; and equilibrated with 3 mL of 0.1% trifluoroacetic acid. Then peptide samples were loaded into the cartridges, washed with 3 mL of 0.1% trifluoroacetic acid and 1 mL of 0.5% acetic acid, and then sequentially eluted first with 0.5mL of 40% acetonitrile, 0.5% acetic acid and then with 0.5 mL of 70% acetonitrile, 0.25% acetic acid. 20 μg and 200 μg aliquots of eluted peptides were dried by vacuum centrifugation and stored at −80°C for proteomic and phosphoproteomic analysis, respectively.

#### Phosphopeptide enrichment

Phosphopeptide enrichment was done by immobilized metal affinity chromatography (IMAC). 200 μg of peptides were resuspended in 150 μl 80% acetonitrile, 0.1% trifluoroacetic acid. To prepare IMAC slurry, Ni-NTA magnetic agarose (Qiagen) was stripped with 40 mM EDTA for 30 min, reloaded with 10 mM FeCl_3_ for 30 min, washed 3 times and resuspended in 80% acetonitrile, 0.1% trifluoroacetic acid. Phosphopeptide enrichment was performed using a KingFisher Flex robot (Thermo Scientific) programmed to incubate peptides with 150 μl 5% bead slurry for 30 min, wash 3 times with 150 μl 80% acetonitrile, 0.1% trifluoroacetic acid, and elute with 60 μl 1:1 acetonitrile:1% ammonium hydroxide. The eluates were acidified with 30 μl 10% formic acid, 75% acetonitrile, dried by vacuum centrifugation, and stored at −80°C until mass spectrometry analysis. Both sample processing and phosphopeptide enrichment were performed in technical duplicate.

#### LC-MS/MS analysis

Peptide and phosphopeptide samples were dissolved in 4% formic acid, 3% acetonitrile, loaded onto a 100 μm ID x 3 cm precolumn packed with Reprosil C18 3 μm beads (Dr. Maisch GmbH), and separated by reverse phase chromatography on a 100 μm ID x 30 cm analytical column packed with 1.9 μm beads of the same material and housed into a column heater set at 50°C. As peptides eluted off the column they were online analyzed by mass spectrometry.

Peptides for proteome analysis were eluted into a Q-Exactive (Thermo Fisher) mass spectrometer by gradient elution delivered by an EasyII nanoLC system (Thermo Fisher). The gradient was 9-30% acetonitrile in 0.125% formic acid over the course of 90 min. The total duration of the method, including column wash and equilibration was 120 min. All MS spectra were acquired on the orbitrap mass analyzer and stored in centroid mode. Full MS scans were acquired from 300 to 1500 m/z at 70,000 FWHM resolution with fill target of 3E6 ions and maximum injection time of 100 ms. The 20 most abundant ions on the full MS scan were selected for fragmentation using 2 m/z precursor isolation window and beam-type collisional-activation dissociation (HCD) with 26% normalized collision energy. MS/MS spectra were collected at 17,500 FWHM resolution with fill target of 5E4 ions and maximum injection time of 50 ms. Fragmented precursors were dynamically excluded from selection for 30 s.

Phosphopeptides for phosphoproteome analysis were eluted into a Velos Orbitrap (Thermo Fisher) mass spectrometer by gradient elution delivered by an Easy1000 nanoLC system (Thermo Fisher). The gradient was 9-23% acetonitrile in 0.125% formic acid over the course of 90 min. The total duration of the method, including column wash and equilibration was 120 min. Full MS scans were acquired in the orbitrap mass analyzer and recorded in centroid mode. Mass range was 300 to 1500, resolution 60,000 FWHM, fill target 3E6 ions, and maximum injection time 100 ms. Each MS scan was followed by up to 20 data-dependent MS/MS scans on the top 20 most intense precursor ions with 2 m/z isolation window, collision-induced dissociation (CID) with 35% normalized collision energy, and acquired on the ion trap. MS/MS spectra were collected at 17,500 FWHM resolution with fill target of 5E4 ions and maximum injection time of 50 ms. Fragmented precursors were dynamically excluded from selection for 30 s.

#### MS data analysis

Acquired mass spectra raw files were searched with MaxQuant v1.6.0.1 (*28*) against the Uniprot mouse canonical plus isoform protein sequence database (downloaded August 2017, 51434 entries) with common contaminants added. The precursor mass tolerance was set to 6 ppm, and the fragment ion tolerance was set to 20 ppm. A static modification on cysteine residues and variable modifications of methionine oxidation and protein N-terminal acetylation were used for all searches except for phosphopeptide-enriched samples where phosphorylation on serine, threonine and tyrosine residues was also included as variable modification. Trypsin/P was the specified enzyme allowing for up to two missed cleavages. The minimum required peptide length was seven residues. The target-decoy database search strategy was used to guide filtering and estimate false discovery rates (FDR). All data were filtered to 1% FDR at both peptide and protein levels. The ‘‘match between runs’’ option was enabled with a time window of 0.7 min to match identifications between replicates. Proteins with at least two peptides (one of them unique to the protein) were considered identified. Label free quantification (LFQ) and intensity-based absolute quantification (iBAQ) algorithms for protein quantification were selected and used for individual protein comparisons between experimental groups and protein organellar ratio calculations, respectively. To determine relative molar abundances iBAQ intensities were further normalized to relative iBAQ values by dividing each protein abundance by the sum of all protein abundances.

For protein and phosphorylation site analysis we used the generated ‘proteinGroups.txt’ and ‘Phospho(STY)Sites.txt’ tables, respectively, after filtering off contaminants and reverse hits. Protein intensity measurements from technical replicates were aggregated, whereas phosphosite intensities were treated individually due to the stochasticity inherent to data dependent mass-spectrometry sampling of individual phosphopeptides. Perseus v1.6.0.7 software (*29*) was used for bioinformatic and statistical analysis using log_2_ transformed data from LFQ normalized protein intensities and median normalized phosphopeptide intensities. Phosphosites were considered localized with a localization probability > 0.75. Annotations were extracted from Gene Ontology (GO) and the Kyoto Encyclopedia of Genes and Genomes (KEGG). Kinase-substrate relationships and kinase-motif sequence information were obtained from the PhosphositePlus database (*30*). Motif extractor tool (motif-x) was used to extract overrepresented motif patterns in sequences surrounding phosphosites (± 7 amino acids) (*31*). Kinase activity was determined using kinase set enrichment analysis (KSEA) which uses Kolmogorov–Smirnov test to assess whether a predefined set of kinase substrates is statistically enriched in phosphosites that are at the two of extremes of a ranked list defined by their differential regulation (*21*). KSEA activity denotes the log_10_-transformed p-values from the enrichment analysis of kinase substrates and is signed based on the average sign of all substrates (i.e. if the majority of substrates present increased or reduced phosphorylation, the kinase is predicted as activated (+) or inactivated (-), respectively).

As a proxy of mitochondrial size, we used the ratio between the summed abundances of all mitochondrial proteins and the summed abundances of outer mitochondrial membrane proteins, as classified by Mitocarta2.0 (*32*) and GO cellular component (GOCC) annotation data. Similarly, for other organelles (Golgi apparatus and endoplasmic reticulum) we used the ratios between total organelle and organelle membrane summed abundances using annotation data from GOCC. These organellar size calculations rely on the principle that changes in the ratio of proteins between the surface and the interior of the organelle would reflect changes in their surface to volume relationship, and hence in their size; and on the assumption that subcellular protein concentrations remain constant. Using a similar rationale we assessed changes in mitochondrial cristae volume by measuring the ratios of summed abundances between mitochondrial intermembrane space and matrix proteins according to the annotation provided by recent proteomic mapping studies using mitochondrial specific proximity-tagging assays (*33, 34*). To validate the accuracy of these measurements we compared the ratio of summed abundances between total and plasma membrane proteins in different tissues and cell lines for which their proteome was measured (*35, 36*) and their cell size obtained from literature (Fig. S4).

#### Sample preparation for Western blot

Vehicle-treated *Ndufs4*^*+/+*^ and *Ndufs4*^*-/-*^ mice, or rapamycin-treated *Ndufs4*^*-/-*^ mice were treated daily from P10 until P29. Mice were then fasted overnight (for 12 h), treated with vehicle or small molecule, and re-fed. Unanesthetized mice were then euthanized by cervical dislocation 4.5 hours later. Brains were immediately isolated and flash frozen in liquid N_2_ for Western blot. Brains were ground using a cryogenic homogenizer. Approximately 50 mg of homogenized brain tissue was lysed with 1 mL RIPA buffer containing Pierce™ protease and phosphatase inhibitor tablets (Thermo Fisher) for 30 minutes. Samples were then centrifuged to remove insoluble cell debris, quantified by BCA, diluted to 2 mg/mL, and used for Western blot.

#### Western blotting

Relative protein levels were determined through Western blotting of 20 µg protein lysate diluted in 4X Laemmli Sample Buffer, 10X Reducing Agent, and RIPA buffer. The samples were heated at 95 °C for 5 minutes then were loaded onto a NuPage™ 4-12% Bis-Tris MIDI gel and run at 120V in MOPS running buffer. Protein was transferred to a PVDF membrane using a Trans-Blot^®^ Turbo Transfer System according to manufacturer instructions (an additional 0.2% SDS was added to the transfer buffer). Unless otherwise noted the blots were blocked with 5% BSA or 5% milk in TBST at room temperature for 1-2 hours and incubated overnight at 4°C with primary antibody. Primary antibodies used were: p-S235/236-S6 ribosomal protein (1:1,000, Cell Signaling Technology 4858), S6 ribosomal protein (1:1,000, Cell Signaling Technology 2217), pS473-Akt (1:1,000, Cell Signaling Technology 4060), pT308-Akt (1:500, Cell Signaling Technology 9275), pT450-Akt (1:1,000, Cell Signaling Technology 9267), Akt (1:1,000, Cell Signaling Technology 4691), pS222-PKC-α (1:2,000, ABClonal AP0559), PKC-α (1:2,000, Cell Signaling Technology 2056), pT638/641-PKC-α/β (1:5,000, Cell Signaling Technology 9375), PKC-β (1:500, Cell Signaling Technology 46809), pT674-PKC-γ (1:1,000, Abcam ab5797), PKC-γ (1:2,000, Santa Cruz Biotechnology sc-166385), pS176/180-IKKα/β Antibody II (1:1000, Cell Signaling Technology 2697), IKKα (1:200 in 1% milk, Novus Biologicals NB-100-56704), pS32-IκB (1:500, Santa Cruz Biotechnology sc-8404), IκB (1:50,000 in 5% milk, Abcam ab-32518), or β-actin HRP conjugate (1:25,000 at room temperature for 2 hours, Cell Signaling Technology 5125). The blots were then rinsed three times at room temperature with TBST for 10 minutes each. The blots were incubated with secondary antibody in 5% BSA (or 5% milk for IκB and IKKα) for 1-2 hours. Secondary antibodies used were donkey anti-rabbit IgG HRP conjugate (1:10,000 or 1:25,000, Thermo Scientific 31458) or mouse-IgGκ BP HRP conjugate (1:2,000, Santa Cruz Biotechnology sc-516102). Blots were thoroughly rinsed with TBST and developed after incubation with Amersham™ ECL™ Western Blotting Detection Reagent for 5 minutes.

### Supplementary Text

#### Rapamycin-associated changes in brain proteome and phosphoproteome of wild-type mice

To determine whether the rapamycin associated changes in brain proteome and phosphoproteome were specific to the Ndufs4 KO model we assessed the effect of the same rapamycin treatment in wild-type mice (Fig. S8A, Table S4, Table S5). Wild-type mice treated with rapamycin also showed reduced body and brain weight (Fig. S8B, S8C). Furthermore, rapamycin associated changes in the proteome and phosphoproteome between wild-type and knock-out mice were strongly correlated (Fig. S8D, S8E), with mTOR and PKC isoforms especially downregulated with rapamycin in both cases. Reduced levels of mTOR and PKC-γ have also been observed in the hippocampus of rapamycin-treated Tsc1+/-mice (*37*). 2D enrichment analysis of GO cellular component terms in the proteome and phosphoproteome datasets (Fig. S8F, S8G) showed that all significantly enriched terms changed similarly in response to rapamycin treatment in both wild-type and Ndufs4 KO mice except for the NADH dehydrogenase complex (i.e. Complex I). Complex I increased slightly in knock-out but not in wild-type in response to rapamycin (Fig. S8D, S8F). We do not know the causes or mechanisms for this selective control, but hypothesize that other complexes in the respiratory chain may impose an abundance threshold for complex I. Together, these results demonstrate that rapamycin elicits a core response in brain that is mostly independent of mitochondrial disease.

#### Rapamycin effect in intracellular calcium regulation

PKC, PKA and CAMK2 regulate the opening of inositol-triphosphate receptor (Itpr) and ryanodine receptor (Ryr), the most abundant calcium-release channels at the ER membrane (*38, 39*). Thus, we reasoned that rapamycin-mediated changes in PKC, PKA and CAMK2 activities could have consequences in calcium signaling and homeostasis in Ndufs4 KO mice brain. Indeed, rapamycin treatment dramatically reduced the levels of Itpr1 (Fig. S11A), the main ER calcium-release channel in brain, and reversed its hyperphosphorylation at activating sites (Fig. S11B). Rapamycin also reduced phospholipase C (Plcb) abundance, the inositol-trisphosphate generating enzyme (Fig. S11A), while increasing the levels of Itpr1-inhibitor Erp44. Unlike Itpr1, Ryr2 activating sites were hyperphosphorylated upon rapamycin treatment (Fig. S11B) which can be explained by differential activation mechanisms. Itpr1 calcium-release activity is stimulated by PKC and CAMK2 phosphorylation but restricted by PKA (*40*), whereas Ryr2 channel opening is promoted by both PKA and CAMK2 phosphorylation (*41, 42*). In addition, rapamycin can alter the activity of both channels by dissociating its interaction with FBKP (*43, 44*). Overall, these results evidence a tighter control of intracellular calcium homeostasis in KO Ndufs4 mice brains upon rapamycin treatment.

## Supplementary Figures

**Data S1. (separate Excel file)**

Proteins quantified by mass spectrometry in wild-type (WT), Ndufs4 knock-out (KO) and rapamycin treated Ndufs4 knock-out (KR) mouse brains. Log_2_ LFQ normalized intensities, raw intensities and iBAQ normalized intensities are provided.

**Data S2. (separate Excel file)**

Phosphosites quantified by mass spectrometry in wild-type (WT), Ndufs4 knock-out (KO) and rapamycin treated Ndufs4 knock-out (KR) mouse brains. Log_2_ median normalized intensities are provided.

**Data S3. (separate Excel file)**

Proteins quantified by mass spectrometry in wild-type (WT) and rapamycin treated wild-type (WR) mouse brains. Log_2_ LFQ normalized intensities, raw intensities and iBAQ normalized intensities are provided.

**Data S4. (separate Excel file)**

Phosphosites quantified by mass spectrometry in wild-type (WT) and rapamycin treated wild-type (WR) mouse brains. Log_2_ median normalized intensities are provided.

**Fig. S1.**
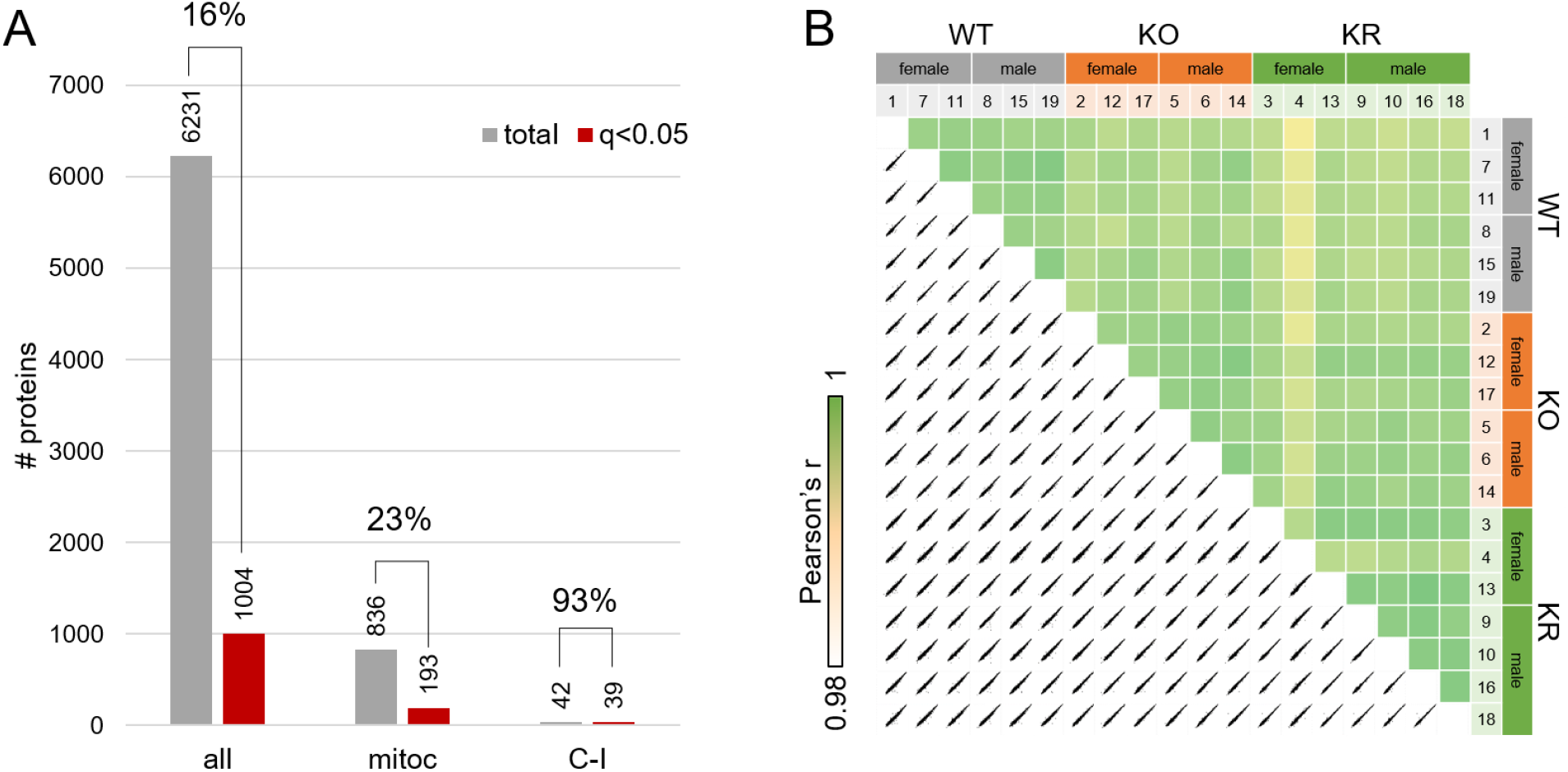
Proteome analysis statistics. (**A**) Number of total (all), mitochondrial (mitoc) and Complex I (C-I) proteins quantified and those with significant changes among experimental groups (ANOVA, FDR q-value < 0.05). (**B**) Correlations of log_2_ transformed LFQ-normalized protein abundance measurements among samples.

**Fig. S2.**
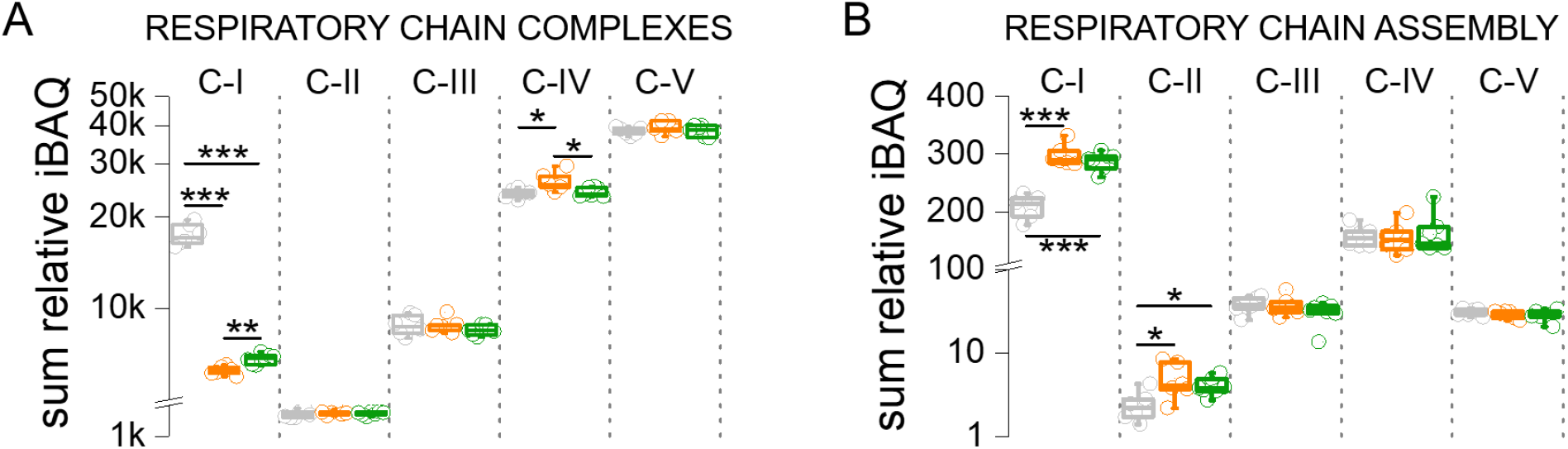
Aggregated protein abundance changes in respiratory chain complexes (A) and respiratory chain assembly proteins (B). Sum of relative iBAQ intensities for all members of each complex or protein group were used.

**Fig. S3.**
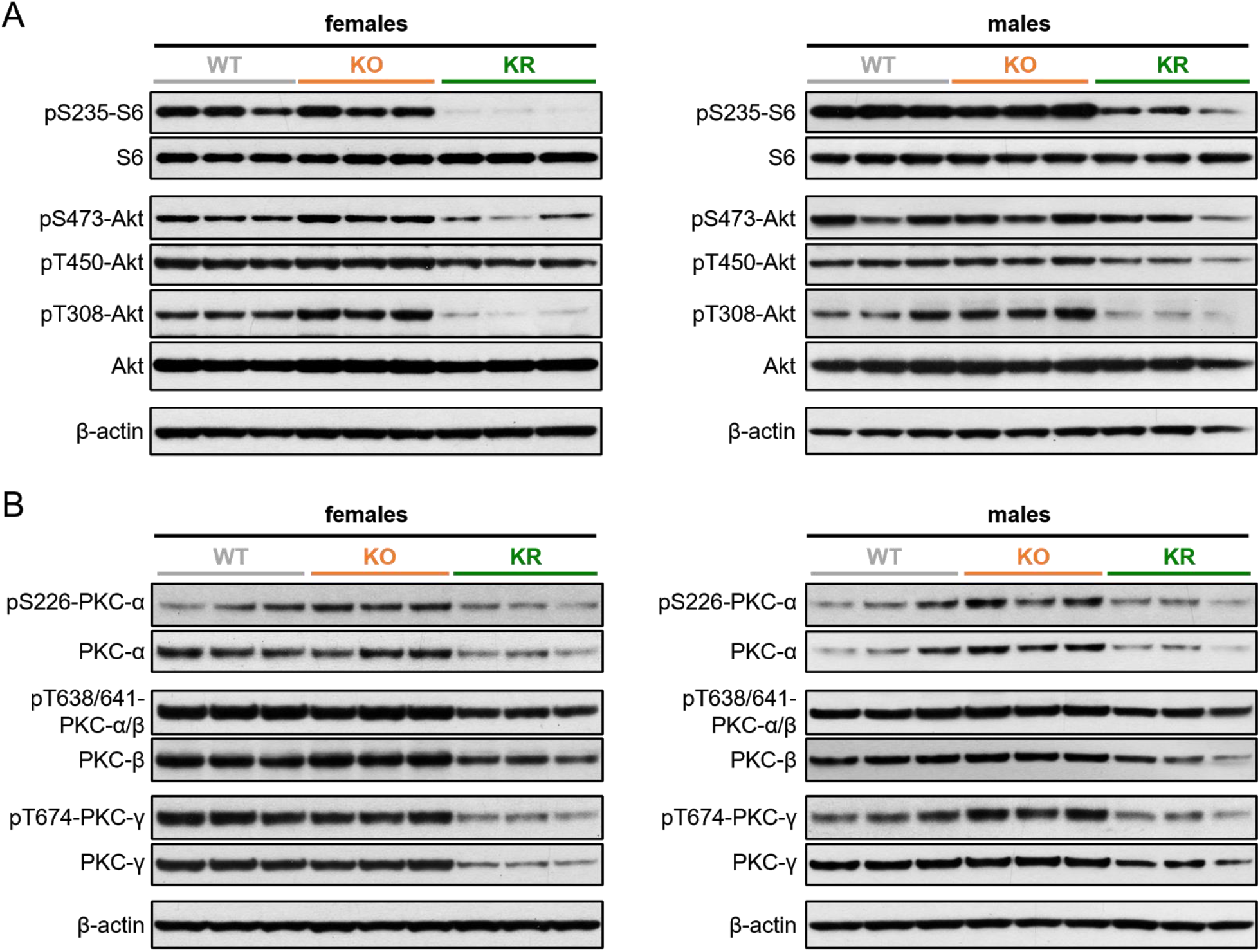
Western blot analysis of mTOR activity. **(A, B)** Western blot analysis of brain lysates from P30 wild type (WT) and Ndufs4 KO mice treated daily with vehicle (KO) or rapamycin (KR) from P10 to P30 suggests that rapamycin treatment led to deactivation of mTORC1 (decreased phospho-S6 and phosphoT308-Akt) and mTORC2 (decreased phosphoS473-Akt and phospho-PKC) **(A)**, as well as decreased phosphorylation and abundance of PKC isoforms **(B)**. Each lane corresponds to a brain lysate from a single mouse.

**Figure S4.**
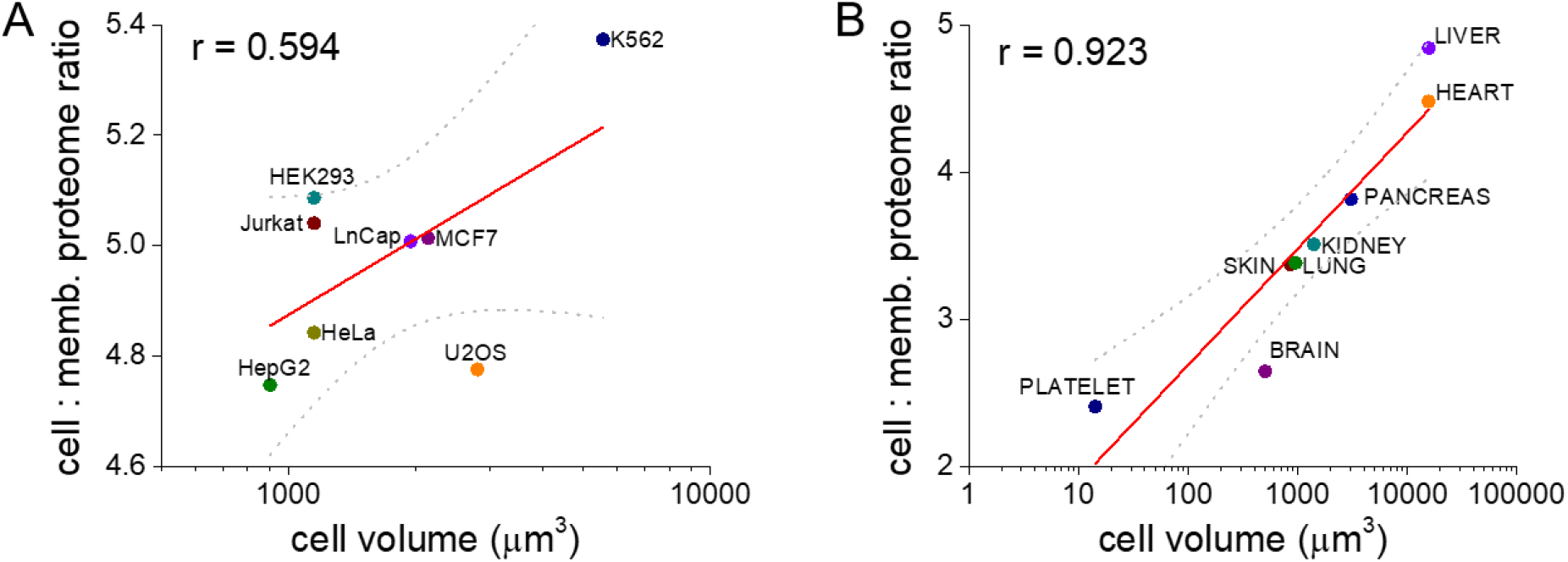
Evaluation of the ratio between whole cell and membrane proteome abundance as an estimate of cell size. (**A**) Correlation of the ratio cell : membrane proteome in human carcinoma cell lines (*35*) and cell volume (data from EMD Millipore cell counter). (**B**) Correlation of the ratio cell : membrane proteome in human tissues (*36*) and tissue specific cell type volumes obtained from the literature. Regression curves are indicated by a straight red line while grey dotted curves indicate 95% confidence interval.

**Fig. S5.**
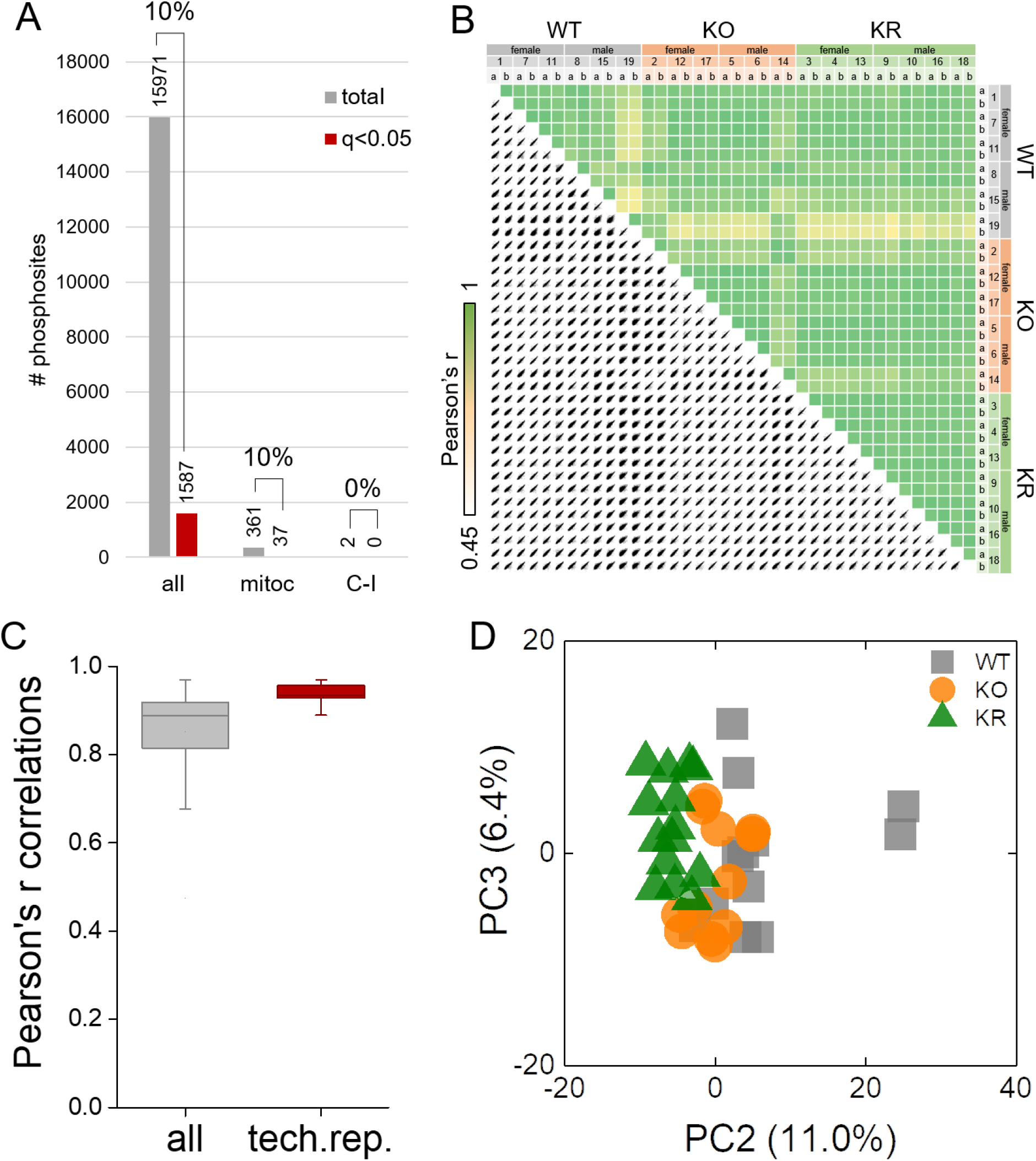
Phosphoproteome analysis statistics. (**A**) Number of total and mitochondrial phosphorylation sites quantified and those with significant changes among experimental groups (ANOVA, FDR q-value < 0.05). (**B**) Correlations of log_2_ transformed median-normalized phosphorylation site intensity values among samples. Two technical replicates of IMAC phosphopeptide enrichment were performed to increase coverage. (**C**) Distribution of Pearson’s r correlation values among all samples and only for technical replicates. (**D**) PCA analysis of log_2_ transformed median-normalized phosphorylation site intensities data.

**Fig. S6.**
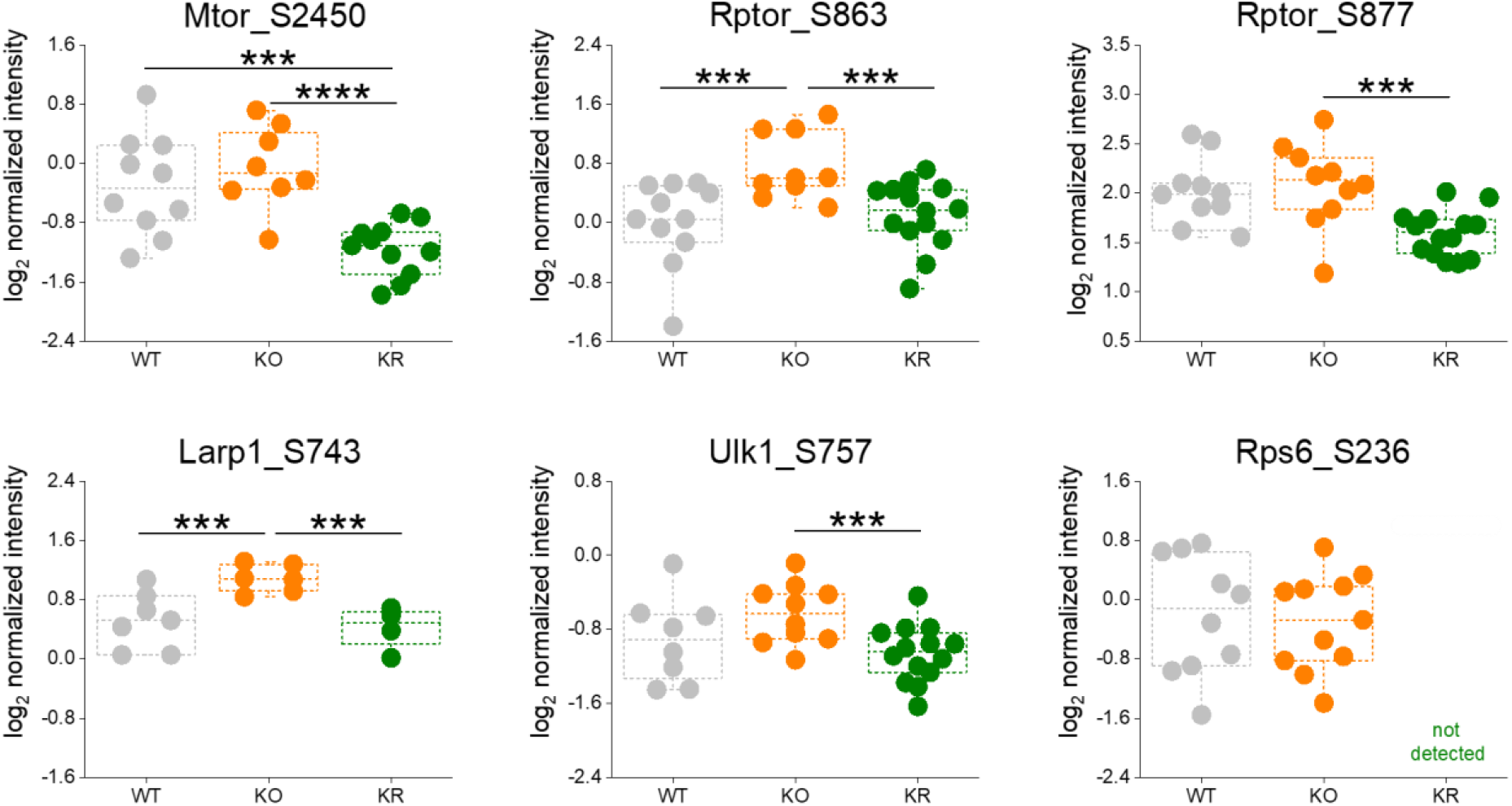
Phosphorylation sites on proteins of the mTOR complexes and associated substrates that show significant changes among experimental groups. T-test significance p-values are indicated (*** p < 0.001; **** p < 0.0001).

**Fig. S7.**
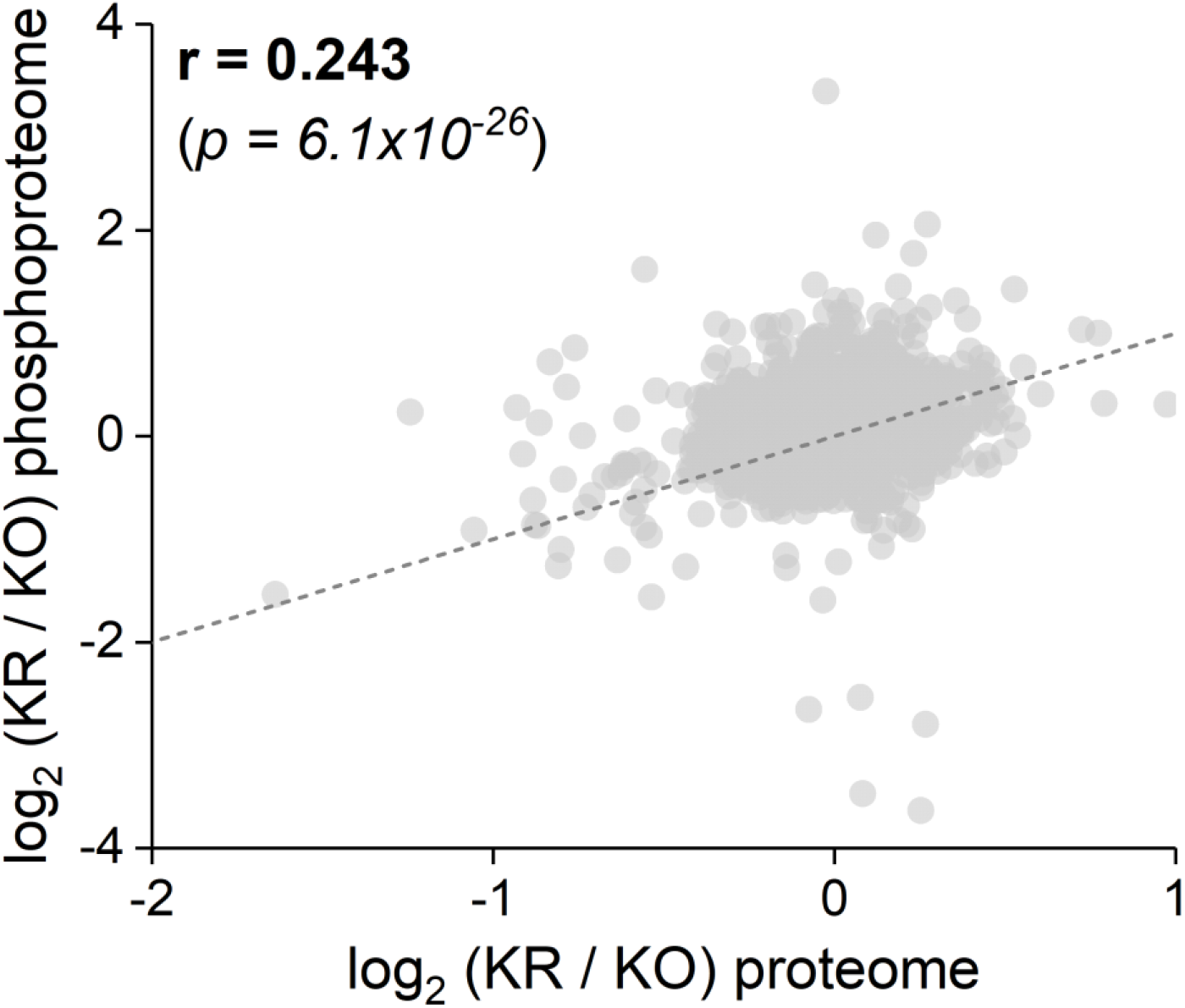
Correlation between rapamycin effects in Ndufs4 KO mice at the proteomic (*x* axis) and phosphoproteomic (*y* axis) levels. Pearson’s r coefficient and goodness-of-fit test p-value are indicated.

**Fig. S8.**
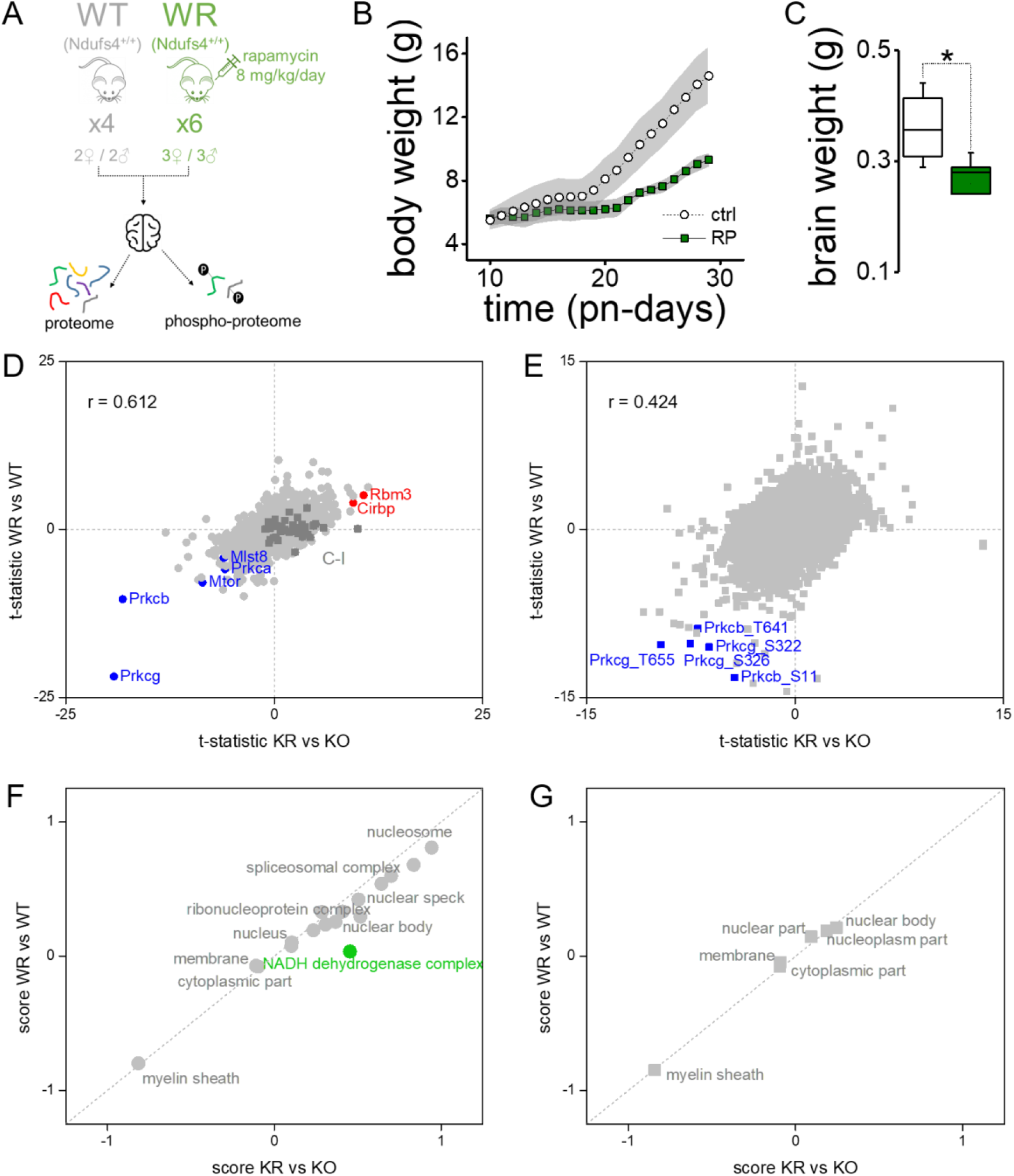
Rapamycin exerts similar effects in brain of wild type mice than Ndufs4 KO mice. (**A**) Experimental design testing rapamycin effect in brain from wild type mice. (**B**) Body weight gain in mice from the two experimental groups. (**C**) Total brain weight at the end of the experimental trial. T-test significance p-values are indicated (* p < 0.05). (**D**,**E**) Comparison of rapamycin mediated changes between wild-type and knock-out mice in individual protein levels (D) and phosphorylation sites (E). Pearson’s correlation values are indicated. (**F**,**G**) 2D annotation enrichment analysis of GO terms comparing the effect of rapamycin between wild-type and knock-out mice in the proteome (F, FDR q-value <0.05) and phosphoproteome (G, p-value < 0.01). Most data points are close to the diagonal dashed line (i.e. identity function), indicating no differences in the effect of rapamycin on wild-type and Ndufs4 KO mice.

**Fig. S9.**
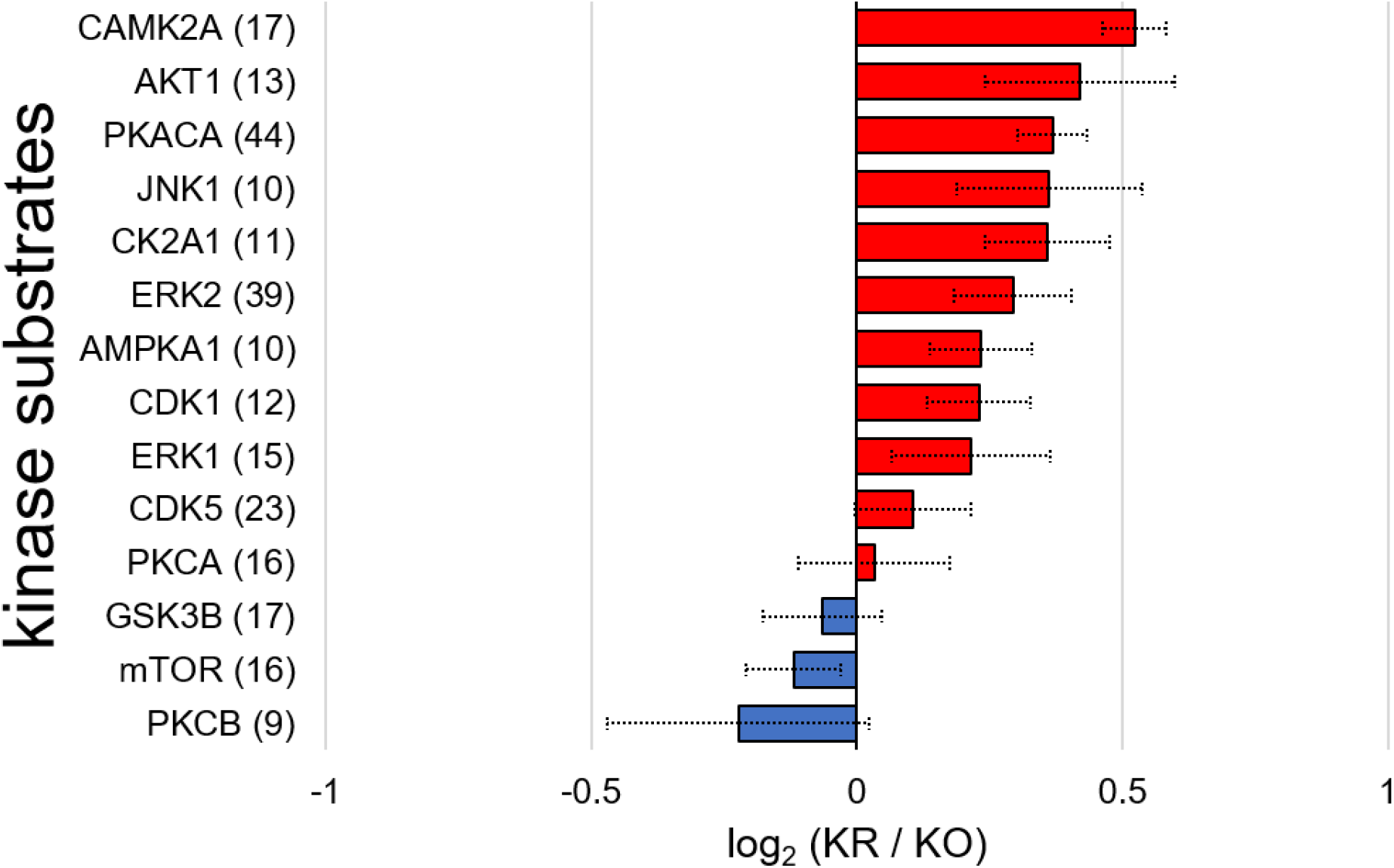
Average (mean ± s.e.m.) changes in phosphorylation of kinase substrates upon rapamycin treatment in brain of Ndufs4 KO mice. Only kinases with more than 9 substrates found are shown.

**Fig. S10.**
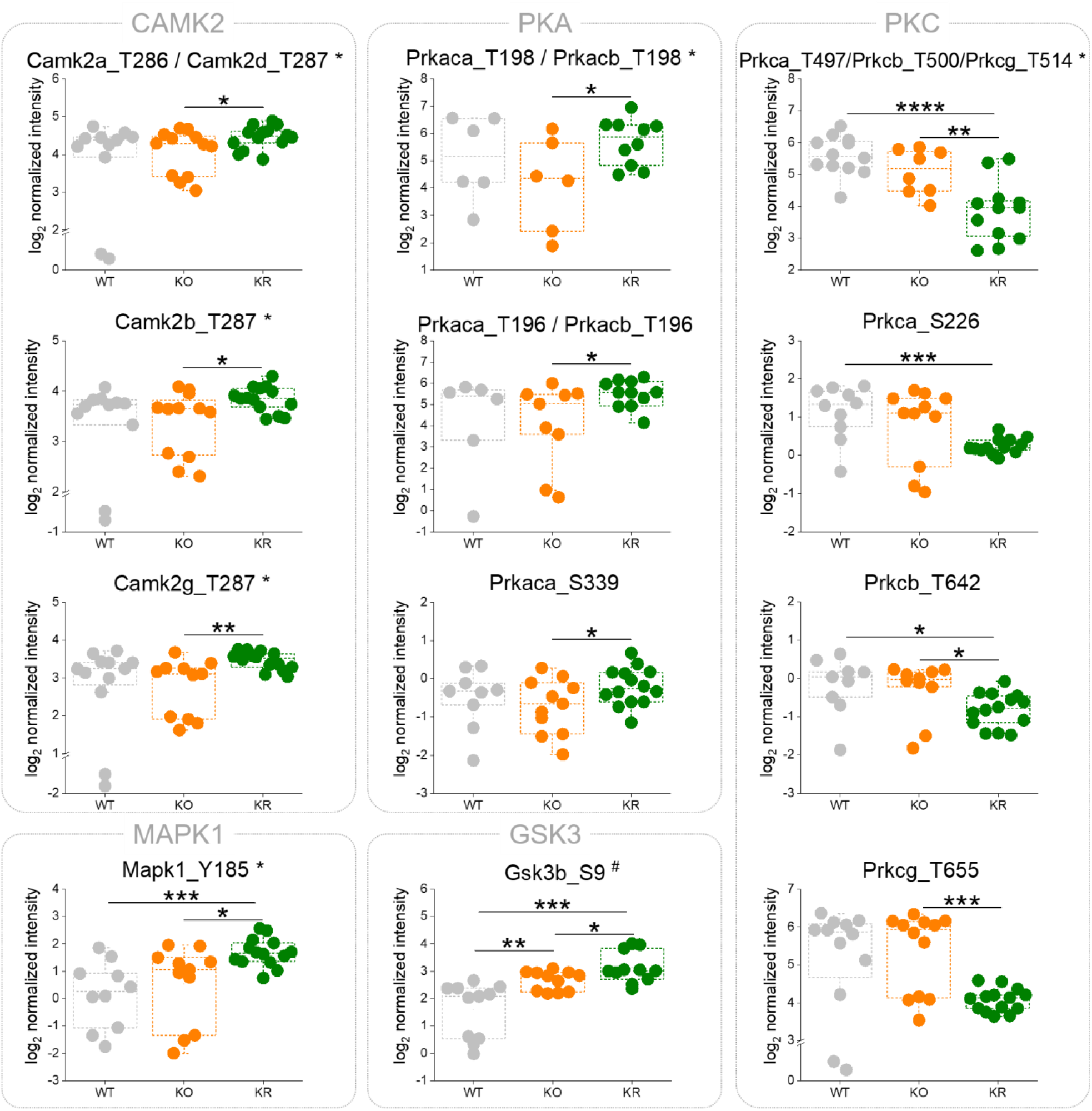
Significant changes in phosphorylation on activity regulatory sites of specified kinases (*activation loop sites, #inhibitory sites). T-test significance p-values are indicated (* p < 0.05; ** p < 0.01; *** p < 0.001; **** p < 0.0001).

**Fig. S11.**
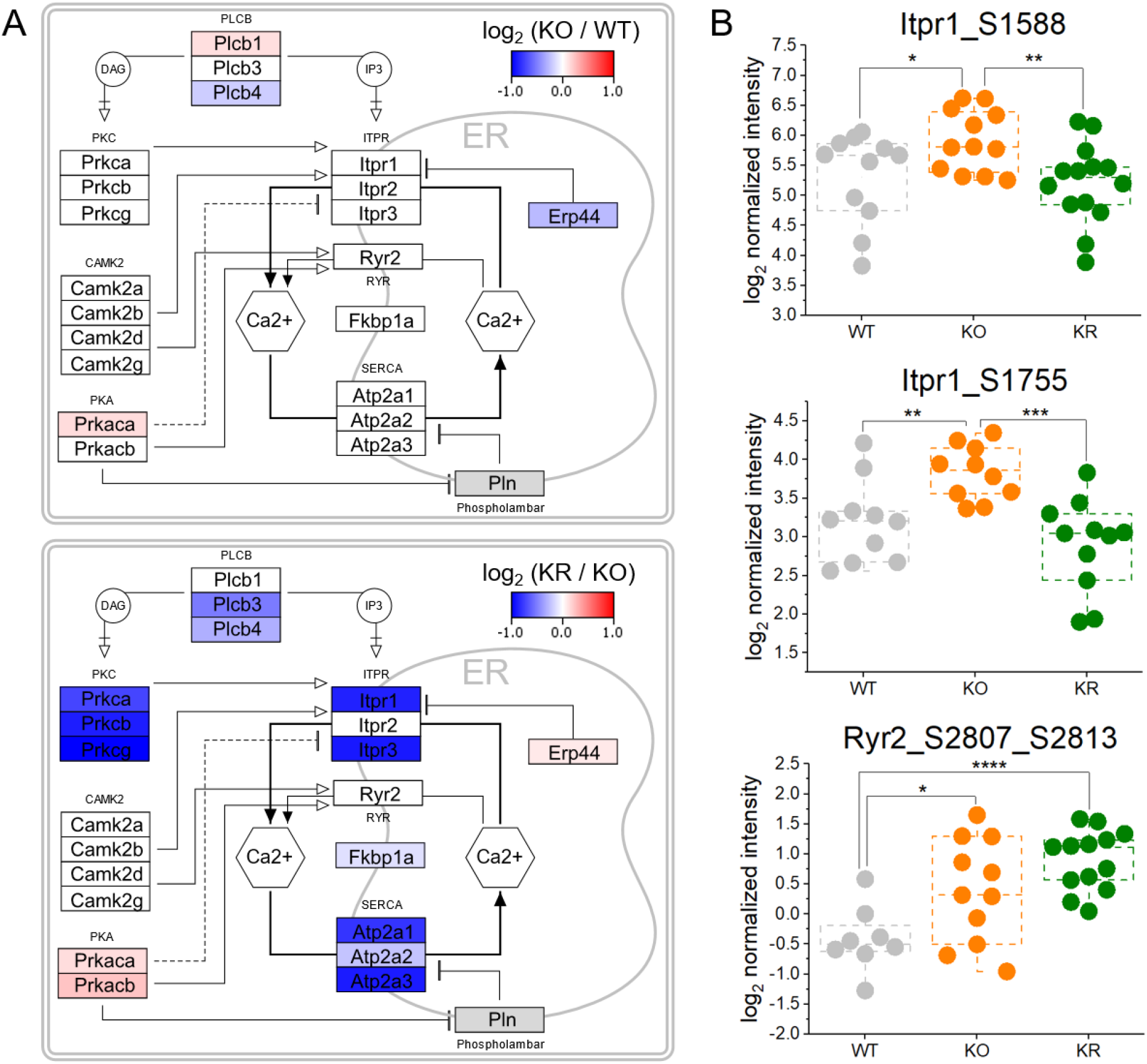
Rapamycin restraints intracellular calcium homeostasis. (**A**) Mapping of protein abundance changes in proteins involved in intracellular calcium homeostasis between the Ndufs4 KO (KO) and wild-type (WT) groups (top panel), and between the Ndufs4 KO mice treated (KR) and untreated (KO) with rapamycin. Only significant changes are colored (t-test p-value > 0.05), non-quantified proteins are shown in grey. (**B**) Significant changes in activating phosphorylation sites of the main two calcium-release channels from the endoplasmic reticulum. T-test significance p-values are indicated (* p < 0.05; ** p < 0.01; *** p < 0.001; **** p < 0.0001).

**Fig. S12.**
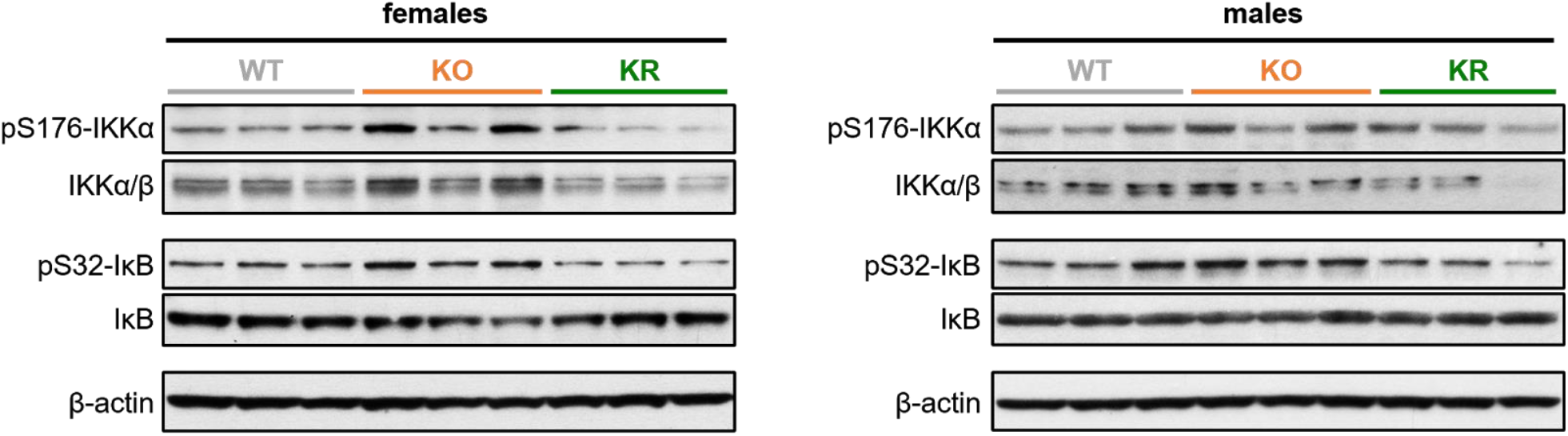
Analysis of NF-κB deactivation by Western blot. Western blot analysis of brain lysates from P30 wild type (WT) and Ndufs4 KO mice treated daily with vehicle (KO) or rapamycin (KR) from P10 to P30 suggests that rapamycin treatment led to deactivation of the NF-κB pathway. Each lane corresponds to a brain lysate from a single mouse.

**Fig. S13.**
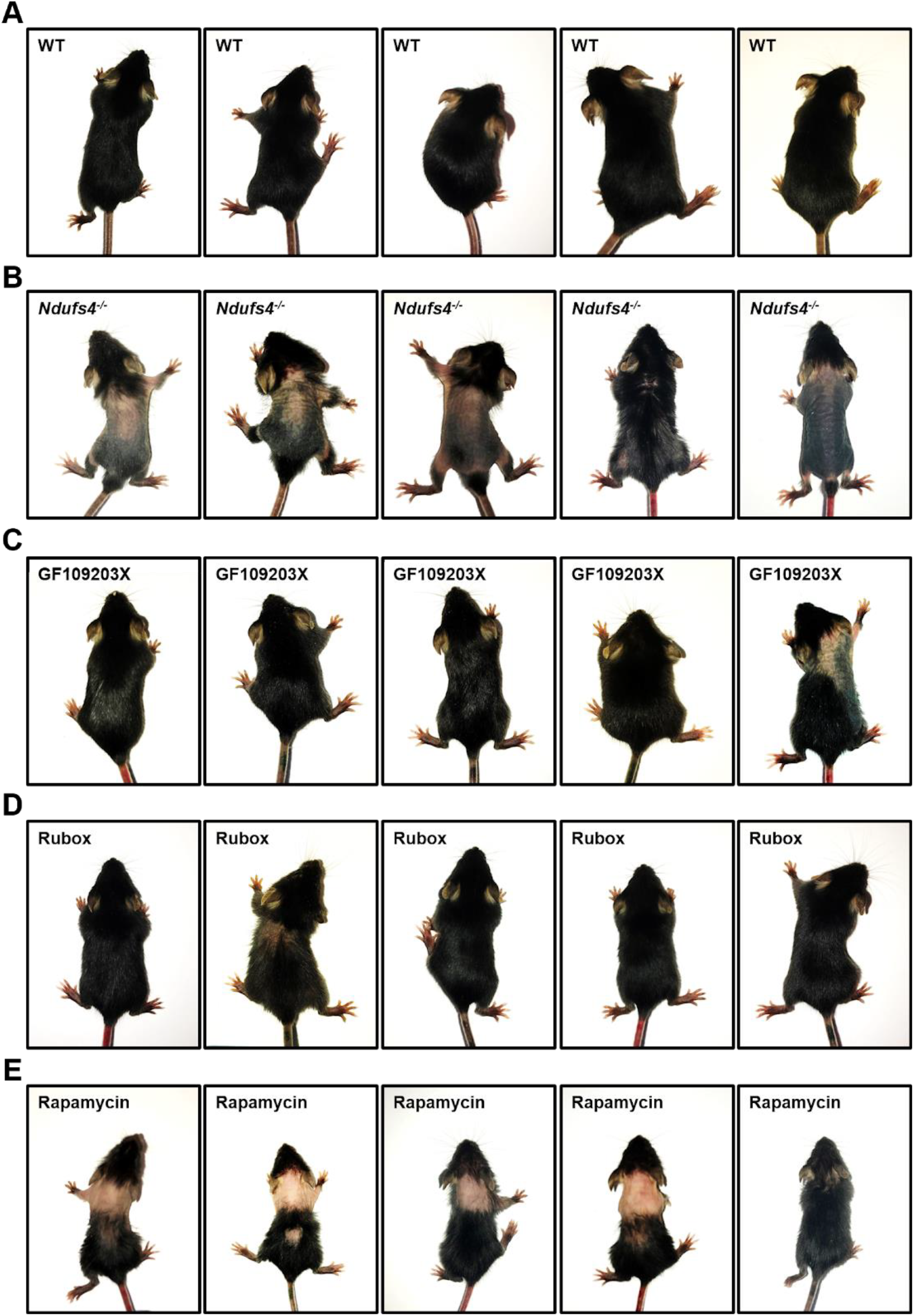
Treatment of Ndufs4 KO mice with PKC inhibitors largely prevents the alopecia phenotype at weaning (∼P21). (**A**) Wild-type mice at weaning show no hair loss. (**B**) Untreated Ndufs4 KO mice normally exhibit alopecia (i.e. hair loss) at 21-days old due to a TLR2/4 innate immune response. In contrast, minimal hair loss was observed in 21-day old Ndufs4 KO mice treated with (**C**) GF109203X and (**D**) ruboxistaurin from P10 to P21. (**E**) Some hair loss was observed in 21-day old Ndufs4 KO mice treated with rapamycin from P10 to P21.

**Fig. S14.**
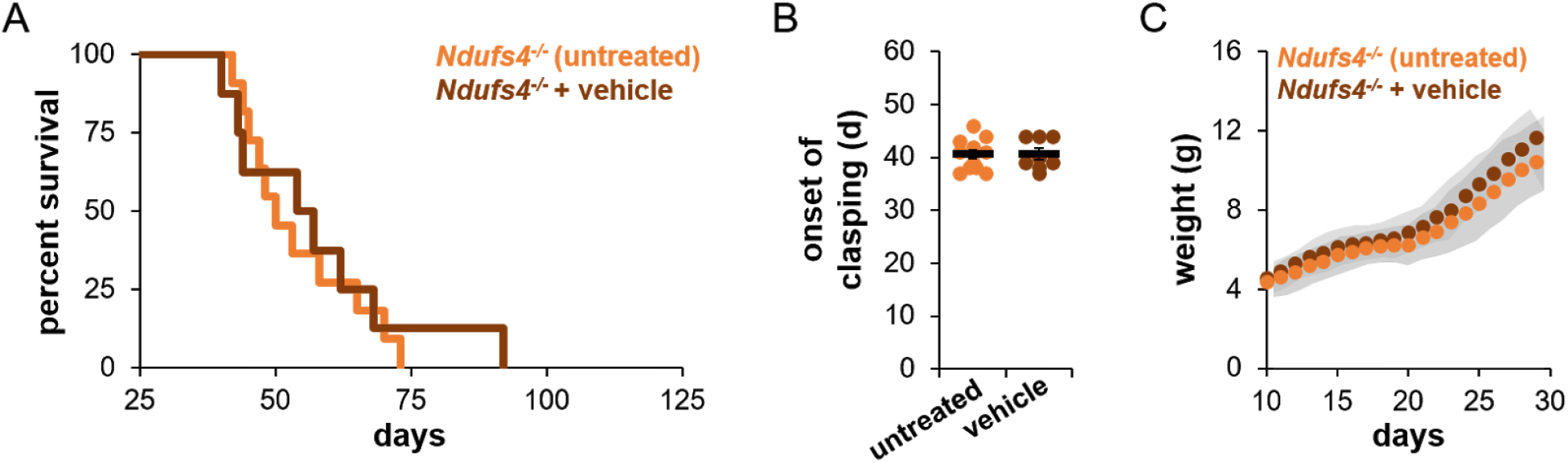
Untreated and vehicle-treated Ndufs4 KO mice exhibit similar symptoms of disease. (**A**) Vehicle treatment does not alter the lifespan of Ndufs4 KO mice compared to untreated controls. Untreated vs. vehicle p-value = 0.7898, log-rank. (**B**) Vehicle treatment does not alter the onset of clasping of Ndufs4 KO mice compared to untreated controls. (**C**) Vehicle treatment does not alter weight or growth of Ndufs4 KO mice compared to untreated controls.

**Table S1.** Mitochondrial proteins with significantly altered phosphorylation sites

| Protein names | Gene name_phosphosites | log<sub>2</sub> *<br>(KO/WT) | t-test<br>q-value<br>KOvsWT | log<sub>2</sub> *<br>(KR/KO) | t-test<br>q-value<br>KRvsKO |
| --- | --- | --- | --- | --- | --- |
| Aflatoxin B1 aldehyde reductase member 2 | Akr7a2_S40 | -0.396 | 0.670 | 1.101 | 0.004 |
| Bola-like protein 1 | Bola1_S81 | -0.213 | 0.819 | 0.793 | 0.007 |
| Glyceraldehyde-3-phosphate dehydrogenase | Gapdh_S146 | 0.173 | 0.754 | -0.471 | 0.020 |
| Glutaredoxin-related protein 5, mitochondrial | Glrx5_S151 | -0.503 | 0.264 | 0.525 | 0.026 |
| Mitochondrial fission factor | Mff_S146 | 0.592 | 0.448 | 0.554 | 0.038 |
| Mitochondrial fission factor | Mff_S151 | -0.253 | 0.913 | -1.267 | 0.011 |
| Oxidation Resistance 1 | Oxr1_S59 | 0.036 | 0.976 | -0.628 | 0.003 |
| Rab11 family-interacting protein 5 | Rab11fip5_S480_S482_S486<sup>#</sup> | 0.360 | 0.826 | -0.709 | 0.019 |
| Rab11 family-interacting protein 5 | Rab11fip5_S693 | -0.125 | 0.950 | 1.078 | 0.011 |
| Saccharopine dehydrogenase-like oxidoreductase | Sccpdh_S217 | 0.019 | 0.993 | -1.239 | 0.001 |
| Monocarboxylate transporter 1 | Slc16a1_S230 | -0.242 | 0.707 | 0.370 | 0.036 |
| Monocarboxylate transporter 1 | Slc16a1_S477 | 0.151 | 0.764 | -0.468 | 0.010 |
| Succinyl-CoA ligase subunit beta, mitochondrial | Sucla2_S281 | -0.760 | 0.020 | -0.084 | 0.631 |

\*Only significant (t-test FDR q-values <0.05) log<sub>2</sub> fold changes are color coded: – +
<sup>#</sup> Phosphosites within the same peptide

